# Trained ILC2 prevent IL-17-associated lung injury during helminth infection through a serotonin-dependent mechanism

**DOI:** 10.1101/2025.11.03.683154

**Authors:** Ulrich Membe Femoe, Fungai Musaigwa, Imani Nicolis, Li-Yin Hung, Camila Napuri, Chinwekele Uzoije, Heather L. Rossi, Cailu Lin, Heidi Winters, Danielle R. Reed, Juan Manuel Inclan-Rico, De’Broski R. Herbert

## Abstract

Type 2 cytokine release promotes wound healing and helminth clearance, but it remains unclear whether group 2 innate lymphocytes (ILC2s) and T-helper 2 cells (T_H_2) cells have functionally distinct roles during anamnestic immunity. This study demonstrates that ILC2 can block re-infection and limit tissue injury caused by the helminth *Nippostrongylus brasiliensis (Nb)*. T_H_2 cells were necessary during initial antigen encounter but dispensable for pathogen clearance and lung repair after ILC2 priming. Upon re-infection, trained ILC2 selectively blocked interleukin (IL)-17+ γδT cell expansion and infection-induced lung injury through an Amphiregulin (*Areg*)-independent mechanism. Trained ILC2s had a distinct metabolic gene expression profile marked by elevated tryptophan hydroxylase 1(*Tph1*) and pulmonary serotonin levels were largely ILC2-dependent. Surprisingly, serotonin prevented IL-17-associated lung hemorrhage irrespective of parasite load. We propose that T_H_2-ILC2 interactions drive pathogen control, but ILC2 distinctly control lung tissue repair through serotonin.

## Introduction

Pulmonary tissue injury caused by host dust mites, fungal products and parasitic helminth infections initiates Type 2 inflammation largely driven by CD4+ T-helper 2 (T_H_2) and group 2 innate lymphoid cells (ILC2)^1^. Both lymphocyte populations secrete Type 2 cytokines like interleukins-(IL-) 4, 5, 9, 13 and Amphiregulin (Areg) that drive a myriad of effector mechanisms subsequent to the release of alarmin cytokines like IL-25 and 33 from hematopoietic and non-hematopoietic cells^2,3^. T_H_2 cells and ILC2 functionally overlap in their secretory profile and while ILC2-derived Areg is considered a distinct driver of tissue repair, the demonstration that Areg-production from CD4+ and γδ+ T cell populations also promotes wound healing raises further question over the distinct role(s) for ILC2 and T_H_2 in an immunocompetent host^4,5^. ILC2-T_H_2 interactions have bi-directional importance and cross-regulatory mechanisms of antigen presentation, co-stimulation, and cytokine-release each influence the functional outcomes of primary Type 2 responses ^6–8^. Moreover, T_H_2 cells were considered solely capable of memory Type 2 responses, but, ILC2 have also been shown to function independently of T and B lymphocytes for recall Type 2 inflammation incited by allergen challenge^9^. Thus, a clear distinction between the functions of ILC2 and T_H_2 cells in the context of pathogen resistance and/or tissue repair remains unclear.

Study of mice infected with the parasitic helminth *Nippostrongylus brasiliensis (Nb)* has yielded foundational aspects of T_H_2 and ILC2 biology ^2,10^. *Nb* third-stage infectious larvae (iL3) travel through skin to enter host circulation, lodge in pulmonary capillaries and break out into alveolar space within 2-3 days, causing hemorrhagic lung injury, neutrophilia, and expansion of IL-17+ γδ T cells (p.i.)^11,12^. Lung egress (3-4 days post-infection (d.p.i.)) and entry into the small bowel is followed by rapid cessation of blood loss and downregulation of IL-17 responses through IL-4Rα-dependent signaling^13,14^. During primary infection, the reparative effects of Trefoil factor 2 and IL-13 resolve lung damage by 7 d.p.i and Type 2 immunity driven by Tuft cells and ILC2 expel adult parasites from the intestine within 9-12 days^11,12,15^. WT mice exhibit strong resistance to re-challenge and eliminate >90% of lung larvae by 3 d.p.i through mechanisms that require involvement of basophils, M2, CD4 and/or ILC2 subsets ^16–18^. IL-33 is necessary for secondary resistance to *Nb* through driving ILC2 expansion and IL-33 also facilitates trained ILC2 development in RAG deficient mice inoculated with allergen ^19–22^. Intestinal ILC2 show enhanced glycolytic and oxidative phosphorylation pathways that mediate proliferation and cytokine release^23,24^ but whether trained ILC2 show metabolic gene expression changes and distinct function(s) in pathogen clearance and/or tissue repair during re-infection is unknown.

This study employed temporal CD4 depletion, selective ILC2 deficiency, adoptive transfer and pharmacological approaches to decipher the roles of T_H_2 vs. ILC2 subsets during anamnestic immunity against *Nb*. CD4 depletion prior to re-challenge had marginal effects, but constitutive CD4 depletion abrogated protective Type 2 immunity. Constitutive ILC2 deficiency impaired lung T_H_2 cell expansion and larval killing during secondary challenge, marked by exacerbated lung hemorrhage and increased IL-17+ γδT cell responses. Surprisingly, Areg only functioned to drive proliferative expansion of T_H_2 cells for parasite clearance but could not reduce lung hemorrhage, IL-17+ γδT cell or neutrophil responses. Trained lung nILC2 and iILC2 subsets elicited by *Nb* rechallenge increased their expression of genes involved in glycolytic and oxphos metabolism and Tryptophan hydroxylase1 (*Tph1*) relative to primary lung ILC2. Critically, our data generated from gain and loss of function strategies demonstrated that lung serotonin production was ILC2-dependent and that serotonin prevented lung hemorrhage and IL-17+ γδT cell responses irrespective of pathogen load. Thus, while T_H_2 cells and ILC2s have interdependent functions upon initial antigen encounter, trained ILC2 distinctly produce serotonin for lung tissue repair and suppression of IL-17 dominant inflammation.

## Results

### CD4^+^ T cells prime ILC2s but are dispensable for recall Type 2 immunity

CD4^+^ T cell depletion throughout the course of infection prevents larval killing upon *Nb* re-challenge^25^, but the cross-regulatory nature of T_H_2 and ILC2 prompted us to experimentally distinguish between the initiation and maintenance phases of immunity. To do this, naive C57BL/6 mice were treated with CD4-depleting (clone GK1.5) or isotype control (clone LFT-2) mAb using either a continuous (day -1 and day 14) or a delayed (day 14 and 21) administration strategy (Fig.1A) In both approaches, mice were s.c. inoculated with 650 iL3 on day 0 followed by anthelminthic treatment with pyrantel pamoate on day 10 post-infection to ensure antigen clearance with re-infection at either 3 days (continuous) or 5 days (delayed) after the last mAb administration (Fig.1A) For each approach, mice were euthanized for analysis of total parasite load (lung and intestine), BAL RBC numbers (proxy for lung hemorrhage), and lung inflammatory cell composition as parasites transitioned from lung to intestine. As expected, continuous α-CD4mAb treatment increased fecal egg output but not the delayed strategy when compared to their respective isotype Ab controls during primary infection (Supplemental Fig. 1A,B). Similarly, continuous CD4 depletion abrogated killing of parasites during re-infection (Fig. 1B,C), but surprisingly there was no impact on parasite killing with delayed CD4 depletion (Fig. 1E,F). Our flow cytometry gating strategy to identify myeloid and lymphoid populations in the lung that used a different anti-CD4 mAb (clone RM4-5) (Supplemental Fig. 1C,D) confirmed that GK1.5 effectively depleted lung tissue CD4^+^T cells (Supplemental Fig. 2A,B).

**Figure 1.**
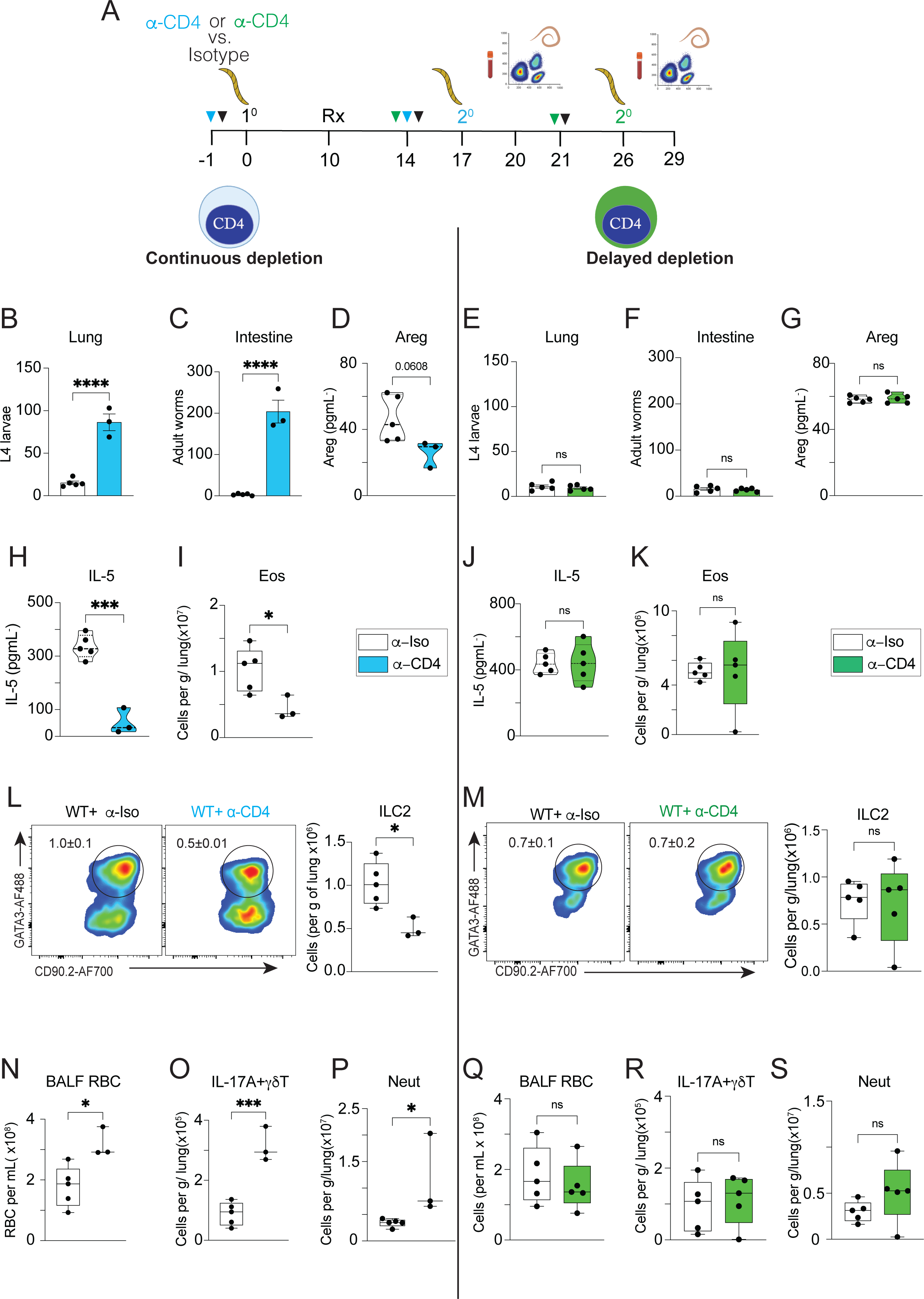
Continuous vs. delayed CD4+T cell depletion changes the outcome of Nb re-infection. **(A)** Schematic showing the continuous vs. delayed CD4 T cell depletion strategy in the context of *Nb* re-infection. Blue indicates continuous strategy. Green indicates delayed strategy. WT C57BL/6J mice (n=3-5 mice/group) were injected (i.p.) with 1mg of anti-CD4 (GK1.5) or isotype mAb one day prior to initial infection with 650 *Nb* infectious larvae (L3) followed by oral gavage with the anthelminthic pyrantel pamoate (Rx) followed by an additional αCD4 or isotype mAb treatment (1mg/mouse), re-infection with 650 *Nb* infectious larvae (L3) and euthanized 3d post-secondary infection. **(B)** Numbers of lung larvae, **(C)** numbers of intestinal worms and **(D)** Areg levels in BAL fluid at 3 days post-re-infection from mice subjected to continuous depletion. **(E)** Numbers of lung larvae, **(F)** numbers of intestinal worms and **(G)** Areg levels in BAL fluid from mice subjected to delayed depletion. **(H)** IL-5 levels in BAL fluid and **(I)** lung eosinophil numbers during continuous depletion. **J**) IL-5 levels in BAL fluid and **(K)** lung eosinophil numbers during delayed depletion **(L)** Representative flow cytometry contour plots and numbers of ILC2 in mice during continuous or **(M)** delayed depletion. **(N)** RBC numbers in BAL fluid, **(O)** and lung tissue numbers of IL-17A+ γδT cells and **(P)** neutrophils during continuous depletion. Q) RBC numbers in BAL fluid, R) and lung tissue numbers of IL-17A+ γδT cells and S) neutrophils in mice during delayed depletion. Cytokine levels were determined by ELISA. Mean and standard error values shown. *P* values were determined by two-tailed Student’s t-tests. *P<0.05, ***P<0.001 ****P<0.0001, ns: non-statistically significant. Representative of two independent experiments.

Bronchoalveolar lavage (BAL) fluid probed by ELISA data revealed that Areg and IL-5 levels tracked with resistance, with significant reductions in the continuous depletion strategy but no change with the delayed strategy (Fig. 1D,H and 1G,J). However, IL-13 levels in BAL were reduced in both approaches (Supplemental Fig. 2C,D). The continuous strategy significantly reduced lung eosinophils and both total ILC2 and IL-5+IL-13+ inflammatory ILC2 subsets (Fig.1I,L; Supplemental Fig. 2E,G). However, delayed CD4 depletion did not reduce eosinophils or total lung ILC2, but moderately reduced IL5+IL-13+ILC2 (Fig.1K,M; Supplemental Fig. 2F,H). Analysis of Arg-1+ M2 macrophage frequencies revealed that both continuous and delayed CD4 depletion reduced this macrophage subset that was implicated in larval killing during *Nb* re-infection (supplemental Fig. 2I,J).

*Nb* larval migration from the vasculature into the alveolar space causes lung hemorrhage associated with neutrophilia and IL-17A secretion by γδT cells^11,12,26,27^. These pathological features were clearly demarcated by CD4 depletion strategy, with continuous CD4 depletion causing significantly increased BAL RBC numbers, lung γδT cell frequencies, IL-17A+ γδT cell numbers and neutrophilia, but delayed CD4 depletion having no impact on these parameters when compared to their respective isotype-treated controls (Fig. 1N-S; Supplemental Fig. 2K-P). These data indicated a temporal requirement for CD4 cells during initial infection in lung ILC2 expansion, parasite killing and control of tissue pathology that was dispensable during re-infection.

### ILC2s are required for protective immunity and tissue repair during *Nb* reinfection

It is well established that ILC2s drive clearance of adult worms from the GI tract during primary *Nb* infection^28–30^, but their importance for secondary resistance is unclear. To test whether anamnestic immunity in the lung required an intact ILC2 compartment, experiments were completed using locus control region 1 deficient mouse strain (*Lcr1*^-/-^) that selectively lacks tissue resident ILC2s^31^. As expected, lung ILC2s were absent at baseline in naïve *Lcr1*^-/-^ mice, but curiously, the lung γδT cell population was significantly elevated relative to littermate controls (Fig. 2A; Supplemental Fig. 3A). *Lcr1*^-/-^ subjected to our *Nb* re-challenge model (Fig. 2B) had increased numbers of parasites in lung and intestine at 3 days post re-challenge compared to WT controls (Fig. 2C,D). While increased fecal egg output during the primary infection phase was expected (Supplemental Fig. 3B), these data indicated an important functional role for ILC2 during *Nb* re-infection.

**Figure 2.**
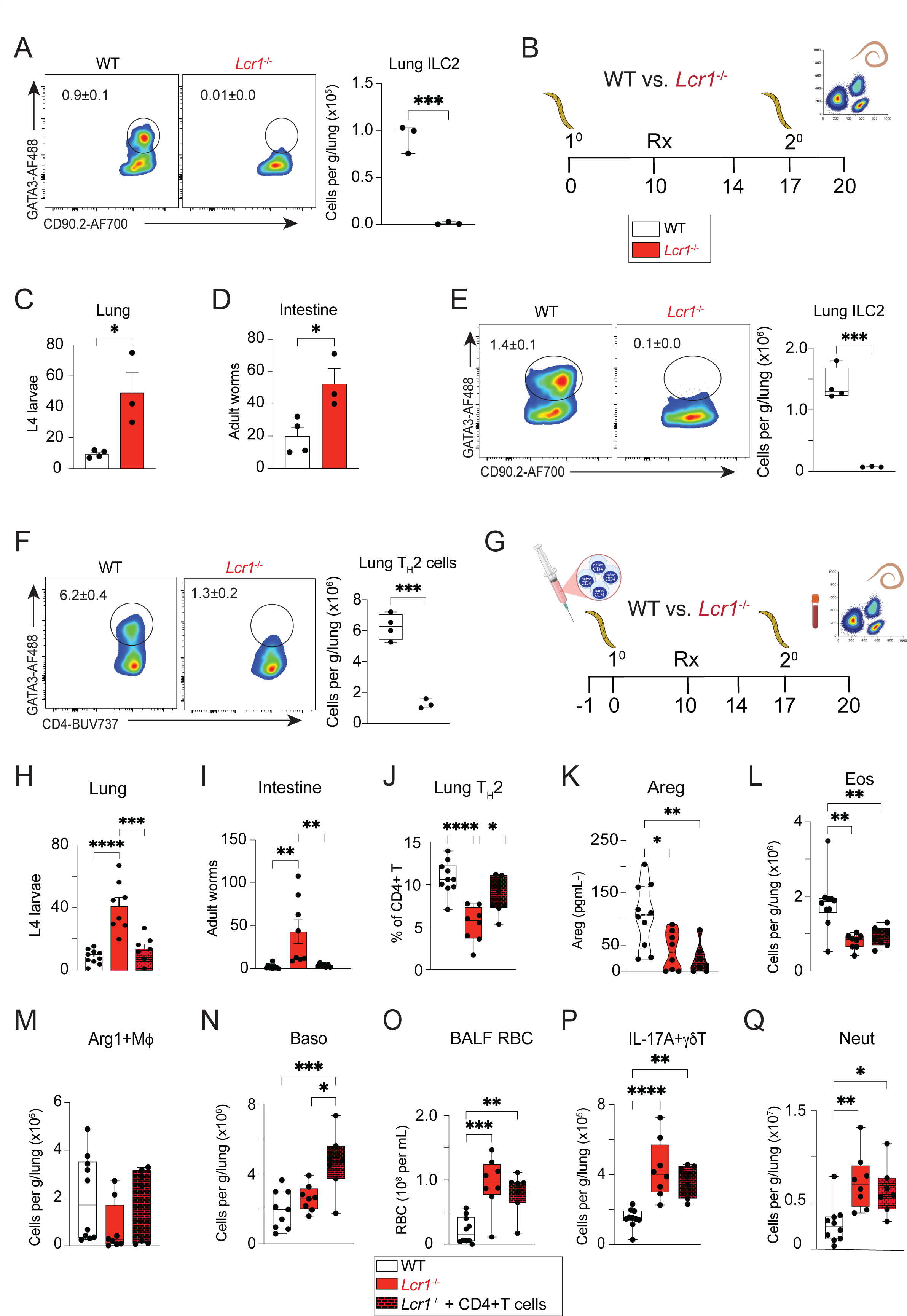
Susceptibility in *Lcr1*^-/-^ mice against *Nb* infection can be reversed by CD4 T cell adoptive transfer, but not hemorrhagic lung injury or IL-17-driven inflammation. **(A)** Representative flow cytometry contour plots showing the total number of lung ILC2 in naïve WT vs. *Lcr1*^-/-^ mice. **(B)** Schematic shows *Nb* re-infection strategy with analysis performed 3 days post-secondary infection. Rx indicates oral gavage with the anthelmintic drug pyrantel pamoate on d10 to ensure clearance of the primary inoculum. **(C)** Numbers of *Nb* larvae recovered from lung tissue and **(D)** intestinal worms recovered at 3 days post-secondary infection. **(E)** Flow cytometry contour plots show total number of lung ILC2 and **(F)** T_H_2 cells in re-infected WT vs *Lcr1*^-/-^ mice 3 days post-secondary *Nb* infection **(G)** Schematic shows strategy for adoptive transfer of CD4+T cells into *Lcr1*^-/-^ mice using 3 million sorted naïve CD4+T cells from naïve WT CD45.1 transferred retro-orbitally into via vein a day prior to primary *Nb* infection. **(H)** Numbers of *Nb* larvae recovered from lung tissue and **I)** intestinal worms recovered at 3 days post-secondary infection in experiments described in “G”. **(J)** Frequency of lung tissue GATA3^+^ T_H_2 cells determined by flow cytometry. **(K)** BAL fluid levels of Areg determined by ELISA. **(L)** Absolute numbers of lung tissue eosinophils, **(M)** Arg-1+M2 macrophages and N) lung basophils in experiments described in “G”. **(O)** Total numbers of red blood cells in the BAL fluid. **(P)** Absolute numbers of lung IL-17A+ γδT cells and **Q)** lung tissue neutrophils as determined by flow cytometry. All data were collected at 3 days post-secondary infection. Mean and standard error values shown. *P* values were determined by two-tailed Student’s t-tests for comparison between two groups or one-way ANOVA followed by Tukey post-hoc test for comparison involving more than 2 groups. *P<0.05, **P<0.01, ***P<0.001, ****P<0.0001, ns: non-statistically significant. these data are pooled from two independent experiments.

### Memory T_H_2 cells fail to control lung injury

However, upon analysis of re-challenged *Lcr1*^-/-^ lung tissue, we noted a combined defect in lung ILC2 numbers and Gata3+T_H_2 cells (Fig. 2E,F). This caveat made it possible that memory T_H_2 cells, if present in sufficient number in *Lcr1*^-/-^ animals, could drive either: 1) pathogen elimination, 2) lung Type 2 cytokine levels, 3) counter regulation of IL-17 driven inflammation, and/or 4) resolution of lung hemorrhage. To address these issues, an adoptive transfer of ∼3x10^6^ sort-purified naïve CD45.1 allogeneic CD4+T cells (CD44^low^CD62L^hi^) was administered to a cohort of *Lcr1*^-/-^ animals and compared to *Lcr1*^-/-^ mock-transferred or WT mice one day prior to initiating the re-challenge protocol with analysis 17 days later (Fig. 2G). Adoptive transfer of naïve CD4+T cells into *Lcr*1^-/-^ mice did not alter clearance rates of the primary infectious inoculum (Supplemental Fig. 3C) but significantly reduced the parasite load in lung and intestine as compared to the mock-treated *Lcr1*^-/-^ cohort and was indistinguishable from the WT group (Fig. 2H,I). Flow cytometry confirmed that the adoptively transferred cells with an effector/memory T_H_2 phenotype (CD45.1^+^CD4^+^CD44^+^GATA3^+^) were recruited to the lung at numbers equivalent to WT re-infected mice (Fig 2J). Areg, IL-5, and IL-13 levels in BAL fluid were not significantly restored by increasing the lung T_H_2 cell population (Fig. 2K; Supplemental Fig. 3D,E) and the numbers of eosinophils and Arg1+M2 macrophages were also not significantly increased (Fig. 2L,M; Supplemental Fig. 3F,G). However, increasing the lung T_H_2 cell population did significantly increase in the basophil population (Fig. 2N; Supplemental Fig. 3H), a cell population consistently shown to drive parasite killing during *Nb* re-challenge ^32,33^.

Surprisingly however, restoring lung T_H_2 cell numbers to WT levels amid ILC2 deficiency was unable to reduce hemorrhagic lung injury caused by migratory larvae as BAL RBC numbers remained significantly elevated (Fig. 2O). In addition, the total lung γδT cell population, IL-17A+ γδT cells and neutrophils remained elevated in both frequency and number at levels equivalent to mock-transferred *Lcr1*^-/-^ animals (Fig. 2P,Q; Supplemental Fig. 3I-K). These data indicated that an intact ILC2 compartment is required for optimal Type 2 cytokine levels, tissue repair, and counter regulation of IL-17-dominant inflammation.

### Primed ILC2s up-regulate *Areg* expression and show gene expression changes indicative of a trained phenotype

Next, the relative abundance of ILC2 and T_H_2 populations were evaluated using confocal microscopy to identify IL-5 expressing cells in IL-5 tdTomato fluorescent reporter mice (i.e. Red5 strain)^34^. Lung IL-5-tdTomato+ cell populations in Red5 mice that were either: naïve, 3 dpi (primary), or 3 dpi (secondary) (Supplemental Fig. 4A-C) were assessed. ILC2s (CD3-Siglec-F-Tdtomato+) were the most abundant IL-5+ population in primary and secondary infection groups followed by T_H_2 cells (CD3^+^Siglec-F^-^Tdtomato^+^) and eosinophils (CD3^-^Siglec-F^+^Tdtomato^+^) (Supplemental Fig. 4D-E). All IL-5-competent cell populations significantly increased during recall responses compared to primary infection (Supplemental Fig. 4E). Flow cytometry was also used to quantify numbers of T_H_2 cells and ILC2 via intracellular staining for GATA3 combined with IL-5 and IL-13. This analysis revealed that while T_H_2 cells outnumbered ILC2 overall, but IL-5^+^ILC2 outnumbered IL-5^+^T_H_2 cells (Supplemental Fig. 4F,G). The number of IL-13^+^ and IL-5^+^/IL-13^+^ T_H_2 cells moderately outnumbered ILC2 subsets with the same cytokine profile (Supplemental Fig. 4H,I).

Natural ILC2 (nILC2) produce large amounts of IL-5, whereas inflammatory ILC2 (iILC2) express both IL-5 and IL-13^18,35,36^, which inspired a hypothesis that perhaps the nILC2 were somehow involved in mitigating lung hemorrhage and γδT cell, IL-17/neutrophil responses. To identify potential candidate genes expressed by ILC2 that could promote tissue repair and immunoregulation, FACS sort-purified ILCs (CD45^+^Lineage^-^CD90.2^+^CD127^+^) from lung tissues of WT C57BL/6 mice at 3 dpi (primary), or 3 dpi (secondary) were subjected to single-cell RNA sequencing (scRNAseq) (Fig. 3A). UMAP (Uniform Manifold Approximation and Projection) plots of this data revealed predominant natural and inflammatory ILC2 clusters with eight minor populations that included: NKT cells, γδ T cells, CD8+ T cells, mast cells, B cells, and cDC (Fig. 3B). As expected, nILC2 and iILC2 cell clusters increased after *Nb* re-challenge (Fig. 3B) and comparison of the top 25 genes in each population confirmed a distinct gene profile Fig. 3C). To assess whether recall infection with *Nb* elicited ILC2 with a trained phenotype, we focused on key metabolic and cytokine receptor genes known to regulate ILC2 function. We found that trained ILC2 elicited upon secondary infection significantly increased expression for *Arg1*, *Cox5a*, *Pkm*, with iILC2 expressing higher levels of *Ki-67*, IL-33R and IL2Ra, but not the IL-7 receptor or GMCSF as compared to nILC2 (Fig. 3D). We postulated that trained ILC2 could drive accelerated tissue repair in *Nb* re-challenged lung tissue^17,37^. Volcano plots of the differentially expressed genes in nILC2 cluster revealed increased (>1.5 FC) in the *Areg* (Fig. 3E). Violin plots of *Areg* expression levels across all ten clusters revealed that nILC2 and iILC2 subsets expressed highest levels (Fig. 3F). We interpreted the data to These data were consistent with trained ILC2 phenotype marked by increased expression of metabolic and tissue reparative genes.

**Figure 3.**
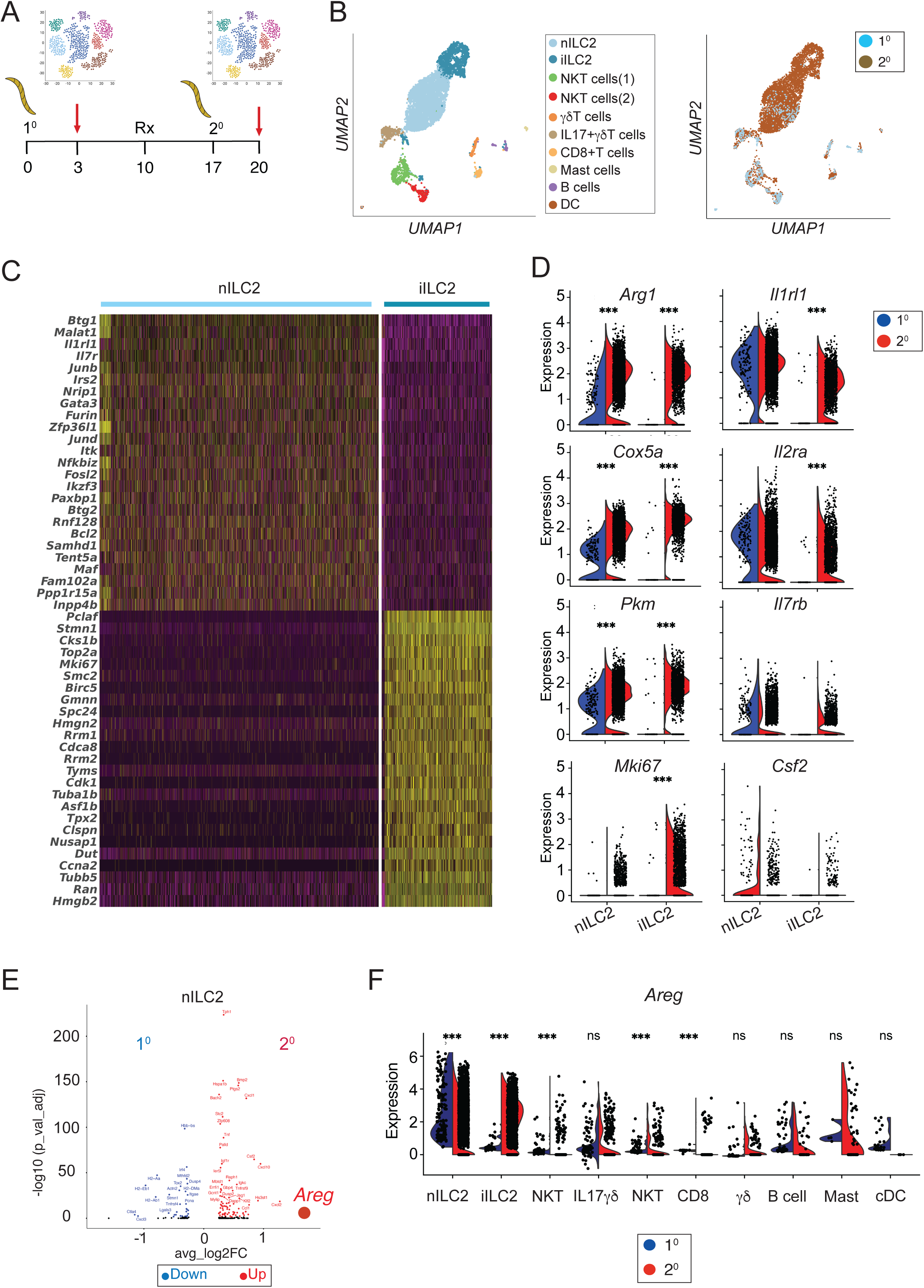
Lung ILC2 subsets show enhanced transcription of metabolic and cytokine receptor genes during *Nb* re-infection. **(A)** Schematic showing strategy for lung ILC2 isolation and single cell RNA sequencing analysis. WT mice (n=20) were infected with 650 iL3 *Nb* and lung tissues were harvested at either 3 days post primary (n=10) or 3 days post-secondary infection (n=10). ILC2 were isolated by MACS using negative enrichment (CD3^-^CD19^-^CD11^-^CD11c^-^CD31^-^CD326^-^ cells) followed by cell sorting to identify ILC2 (Live CD45^+^Lin^-^CD90.2^+^CD127^+^) **(B)** UMAP data shows all cell clusters identified and single cell distribution profile from data combined across primary and secondary infection. **(C)** Heatmap showing transcripts expressed in natural ILC2 (nILC2) vs. inflammatory ILC2 (iILC2) subsets using the FindMarkers function to identify the top 25 differentially expressed genes for heatmap visualization. Expression values were scaled to highlight relative expression patterns. **(D)** Gene expression levels for Arginase1, Cytochrome c oxidase subunit 5a, Pyruvate kinase, Ki-67, IL-1receptor like 1 (ST2), IL-2 receptor alpha, IL-17 receptor B and Colony Stimulating Factor 2 (GM-CSF) compared between natural and inflammatory ILC2 subsets in lung tissues after primary and secondary *Nb* infection. Samples within each Data analyzed using the Wilcoxon–limma model in Seurat’s FindMarkers function. Subset scRNA-seq data retaining nILC2 (3,579 cells) and iILC2 (1,412 cells) from primary (n = 189) and secondary (n = 4,802) samples were used to generate violin plots. Significance was determined using Bonferroni-adjusted p-values across all genes. *** indicates adjusted p-value < 0.001. **(E)** Volcano plots showing differentially expressed genes in the nILC2 cluster in primary vs. secondary responses reveal *Areg* expression in transcripts up-regulated secondary lung nILC2s relative to primary nILC2. The statistics were computed using the Wilcoxon rank-sum test in combination with limma model. The significant threshold was set up as avg_log2FC >0.5 or <-0.5 and the percentage of cells in each group >5% and FDR<=0.05. **(F)** Violin plots showing *Areg* expression levels across all identified cell clusters.

### Amphiregulin drives lung T_H_2 cell expansion and parasite elimination

Various lymphocyte and myeloid populations produce *Areg* for tissue repair in both infectious and non-infectious contexts^38–40^. To investigate whether *Areg* functioned to resolve lung hemorrhage and/or block IL-17 driven inflammation, *Lcr1*^-/-^ mice were inoculated i.n. with 5 ug of rAreg on days 16, 17, and 19 of our re-infection protocol (Fig. 4A). Areg supplementation significantly reduced lung larvae and intestinal worms compared to vehicle-treated controls (Fig. 4B, C). Areg-mediated parasite clearance was accompanied by lung T_H_2 cell expansion, increased BAL levels of IL-5 and IL-13, and increased eosinophil and Arg-1+M2 macrophage numbers in lung tissue (Fig. 4D,E; Supplemental Fig. 5A-D). However, rAreg administration did not reduce BAL RBC numbers or block increased total γδT cell, IL-17A+γδT cell or lung neutrophils compared to the vehicle-treated *Lcr1*^-/-^ cohort (Fig. 4F-H; supplemental Fig. 5E,F). To address whether rAreg had a direct impact on T_H_2 cell biology, FACS-sorted CD4^+^CD44^+^T cells were exposed to rAreg in culture for 48h and analyzed using the self-organizing map (SOM) tool FlowSOM for unsupervised clustering and dimensionality reduction for changes in cytokine/cytokine receptor, transcription factor, and proliferation profiles (Fig. 4I). This analysis revealed 8 clusters that could be stratified by their pattern of IL-5, IL-13, Gata3, ST2, CD44 and Ki-67 expression, with 4 of these clusters significantly increased in frequency due to rAreg exposure. All expressed IL-13 and 3 clusters co-expressed the IL-33R (T1/ST2) and Ki-67 indicating S phase entry (Fig 4J-L, Supplemental Fig. 6A-D). This was corroborated by conventional flow analysis, which showed rAreg exposure significantly increased GATA3^+^T_H_2 cells that expressed Ki-67, IL-5 and IL-13 (Supplemental Fig. 6E-G). Thus, while Areg mediated T_H_2 cell expansion and parasite control it did not augment resolution of lung hemorrhage or block IL-17 responses.

**Figure 4.**
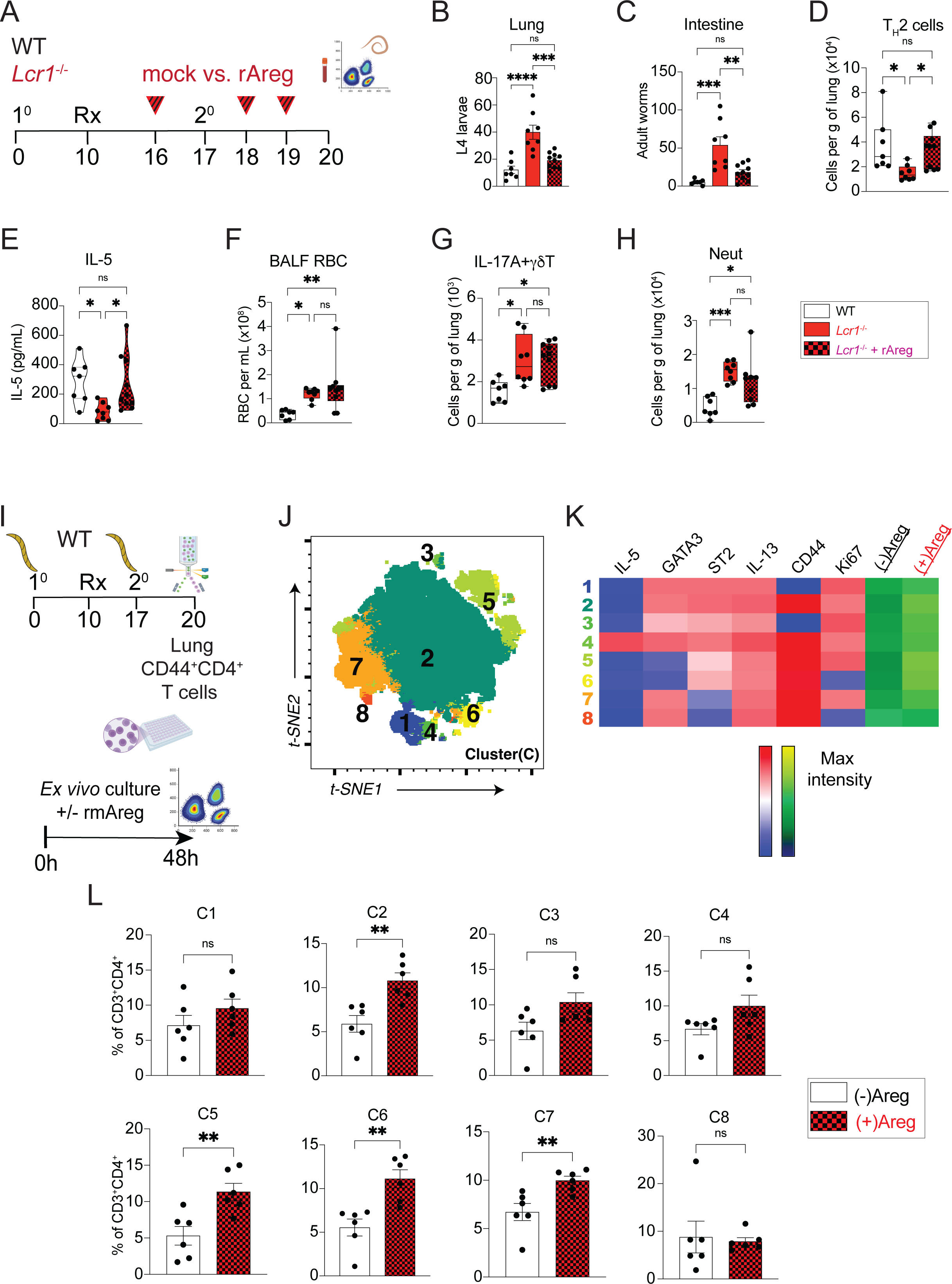
Amphiregulin administration to *Lcr1*^-/-^ mice drives T_H_2 polarization and parasite clearance but does not block lung hemorrhage or IL-17 responses. **(A)** Schematic showing strategy for Amphiregulin supplementation in *Lcr1*^-/-^ mice during re-infection. WT C57BL/6 mice were compared to *Lcr1*^-/-^ or *Lcr1*^-/-^ mice 3 intranasal doses of 5µg of recombinant amphiregulin (rAreg) one day prior and two consecutive days following *Nb* re-infection with 650iL3. **(B)** Numbers of lung larvae and **C)** intestinal worms 3 days post-re-infection. **(D)** Numbers of lung T_H_2 cells determined by flow cytometry. **(E)** IL-5 levels in BAL fluid determined by ELISA, **F)** numbers of RBC in BAL fluid, **G)** number of lung IL-17A+ γδT cells and **H)** lung neutrophils at 3 days post re-infection. **(I)** Schematic showing sort-isolation of antigen-experienced CD4+T cells from Nb re-infected WT C57BL/6 mice followed by 48h *in vitro* culture with rAreg. 300,000 cells were plated with rAreg or vehicle in triplicate wells. **(J)** FlowSOM analysis showing identified clusters expressed via t-SNE dimensionality reduction **(K)** with heatmaps based on their combinatorial expression profile of markers associated with activated T_H_2 cells in experiments described in “I” above. **(L)** Frequency of each FlowSOM clusters 1-8 cells treated with rAreg vs. vehicle. Mean and standard error values shown. *P* values were determined by two-tailed Student’s t-tests for comparison between two groups or one-way ANOVA followed by Tukey post-hoc test for comparison involving more than 2 groups. *P<0.05, **P<0.01, ***P<0.001, ****P<0.0001, ns: non-statistically significant. The figures are the representation of pooled data from two independent experiments.

### Serotonin is necessary and sufficient to control IL-17A associated lung injury during *Nb* reinfection

Upon closer evaluation of trained nILC2 and iILC2 subsets following secondary infection, we noted a moderate increase in tryptophan hydroxylase 1 (*Tph1)* expression that was highly significant (Supplemental Fig. 7A). *Tph1* transcripts were restricted to nILC2 and iILC2 among the 10 identified clusters within our sc-RNA seq data set (Fig. 5A). BAL fluid serotonin levels were measured across experimental protocols used in our study to test whether BAL RBC numbers and IL-17^+^γδ T cell responses were inversely associated with its production. Curiously, serotonin levels were extremely low during 1) continuous CD4^+^T cell depletion, 2) re-challenged *Lcr1*^-/-^ mice, 3) naïve *Lcr1*^-/-^ mice, 4) T_H_2 cell reconstituted *Lcr1*^-/-^ mice or 4) rAreg treated *Lcr1*^-/-^ mice (Fig. 5B). In contrast, BAL serotonin levels were at least 3-fold higher in mice subjected to delayed CD4 T cell depletion protocol that had lung ILC2 numbers equivalent to re-infected WT mice (Fig. 5B). FACS-sorted ILC2s and T_H_2 cells from both primary and secondary infected wild-type (WT) mice were tested for serotonin release in culture. Following stimulation with recombinant IL-33 (rIL-33), ELISA results indicated that only ILC2s produced 5-HT (Supplemental Fig. 7B,C)

**Figure 5.**
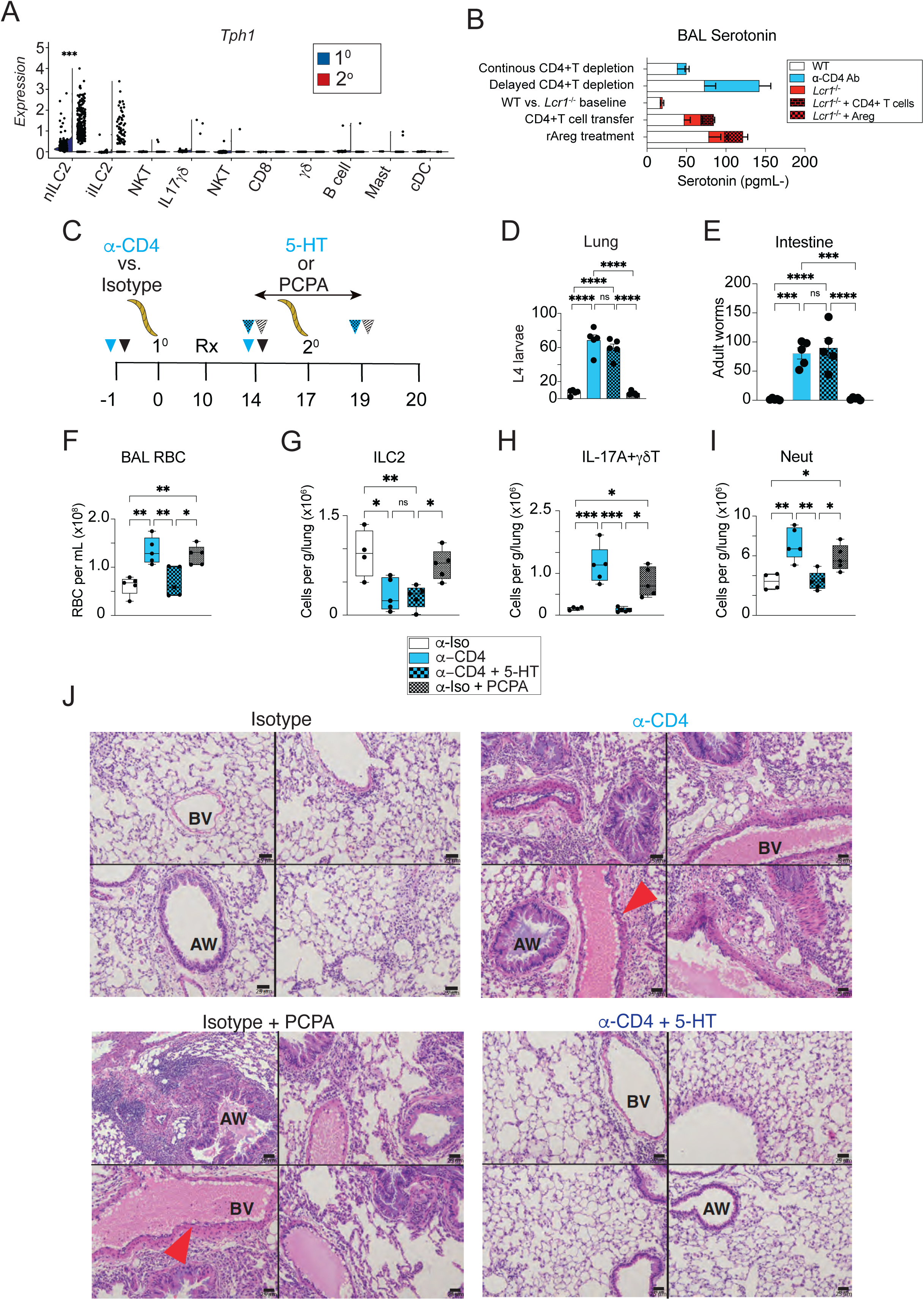
Lung ILC2 subsets are a major source of serotonin(5-HT) that controls Nb infection-induced lung pathology irrespective of parasite burden. **(A)** Violin plots from sc-RNAseq analysis showing *Tryptophan hydroxylase 1* (*Tph1*) expression across all cell clusters identified in “Fig. 3B” reveals significantly increased levels in nILC2 after *Nb* re-infection. **(B)** Levels of serotonin (5-HT) in BAL fluid compared across 5 different experimental approaches employed in this study. **(C)** Schematic showing strategy for 5-HT supplementation vs. Tph1 inhibition in WT C57BL/6 mice subjected to mice continuous CD4 depletion or treated with isotype control, respectively. All mice were subjected to the re-infection approach. Mice subjected to continuous depletion were administered 5-HT (0.1 mg/mL) or vehicle intranasally from day 14-day 19. For Tph1 inhibition, WT mice treated with isotype Ab were intraperitoneally injected with 300 mg/kg of PCPA from day 14-day 19. **(D)** Numbers of lung larvae and **E)** intestinal worms recovered 3 days post-reinfection. **(F)** RBC numbers in BAL fluid **(G)** Numbers of lung ILC2, **(H)** IL-17A+ γδT cells and **(I)** lung neutrophils numbers at 3 days post-reinfection **(J)** Data show **4** representative photomicrographs of H&E stained lung tissue after perfusion at 3 days following reinfection from each of the 4 groups in the experiments described in “C” (scale bar = 25 µm). Red arrows indicate blebbing along endothelial lining. *P* values were determined by one-way ANOVA followed by Tukey post-hoc multiple comparison test. *P<0.05, **P<0.01, ***P<0.001, ****P<0.0001, ns: non-statistically significant. BV= blood vessel. AW= airway. Representative of two independent experiments.

Lastly, two independent, but complementary approaches were used to directly test whether serotonin (5-HT) responsiveness modulated either parasite clearance, lung tissue repair or IL-17 inflammation. The continuous CD4 depletion protocol was employed in WT mice to reduce both lung tissue T_H_2 and ILC2 numbers (Fig. 1L; Supplemental Fig. 2A). As a gain-of-function strategy, WT mice continuously treated with α-CD4mAb were injected daily with 1µg recombinant 5-HT i.n. between d14-19 and evaluated on d20. As a loss-of-function approach, WT mice were administered p-chlorophenyl alanine (PCPA; 300 mg/kg) from d14-19 to inhibit *Tph1*^41^ strictly prior to re-infection (Fig. 5C). Data show no impact on parasite burden upon 5-HT supplementation or Tph1 inhibition (Fig. 5D,E). However, 5-HT treatment significantly reduced RBC numbers whereas PCPA significantly increased BAL RBC numbers despite the absence of lung parasites (Fig. 5F). Total ILC2 numbers were not altered by Tph1 inhibition (Fig. 5G; Supplemental Fig. 7D). 5-HT treatment significantly reduced lung γδT cells,17A+ γδT cells and neutrophils in both frequency and number, but PCPA treatment had the exact opposite effect, with increased IL-17A^+^ γδT cell and neutrophil frequencies (Fig. 5H,I, Supplemental Fig. 7E-G). We surmised that BAL RBC levels were associated with damage around vasculature as worms passed from vasculature into the alveolar space. Indeed, histopathological assessment of formalin fixed paraffin embedded (FFPE) lung tissues subjected to H&E staining revealed that 5-HT-treatment reduced peri-bronchial inflammation and vascular endothelial cell damage in mice with reduced CD4 and ILC2 whereas blocking 5-HT responsiveness with PCPA in re-infected WT mice caused severe exacerbation of inflammatory cell infiltration and vascular injury (Fig. 5J). Collectively, these data indicated that serotonin, (most likely from ILC2) controlled the extent of vascular damage and IL-17A associated lung inflammation during *Nb* re-infection through mechanisms independent of parasite control.

## Discussion

T_H_2 and ILC2 subsets show broad overlap in cytokine profiles, ability to migrate, undergo proliferative expansion, undergo shifts in metabolism and develop memory ^42–44^. Thus, it has remained a puzzling conundrum whether non redundant functions exist for these distinct lymphocyte populations. This study demonstrates that ILC2 distinctly produce the neurotransmitter 5-HT that limits hemorrhagic lung injury and IL-17^+^ γδT cell responses in lung tissues of mice re-infected with the parasitic helminth *Nippostrongylus brasiliensis*. Our work supports the concept that ILC2 undergo innate immune “training” marked by increased expression of genes involved in glycolysis and OXPHOS metabolism and tissue repair *Areg* and *Tph1*. Unexpectedly, Areg was not responsible for pulmonary tissue repair, but rather T_H_2 cell expansion whereas *Tph1*-dependent 5-HT release from ILC2 curtailed hemorrhage and suppressed IL-17+γδT cell/neutrophil responses irrespective of pathogen load. Moreover, our data corroborate reports of extensive cross-regulation between T_H_2-ILC2 populations, as ILC2 deficiency impaired Areg-driven T_H_2 expansion whereas early CD4 depletion impaired ILC2 progression into a trained state. However, once trained, ILC2 could mediate pathogen control, tissue repair and suppression of pathogenic IL-17 inflammation independently of T_H_2 cells. Thus, while interdependent during initial antigen encounter, a division of labor exists between ILC2s and T_H_2 cells for comprehensive recall Type 2 immunity.

Transient pulmonary tissue migration is a conserved feature among many helminth species and migratory *Nb* larvae cause significant damage to vascular and epithelial tissues *en route* to the GI tract^45–47^. Mice infected with *Nb* develop robust host resistance against reinfection and the lung is a key site for parasite attrition driven by type 2 responses that require IL-4, memory T_H_2 cell and STAT6-dependent pathways^17,37,47,48^. Using various models of helminth infection, many have investigated the role of memory T_H_2 cells and shown their importance through IL-4/IL-13 and alternatively activated macrophages (M2) that contribute to parasite elimination, in part via Arg1 ^13,49–51^. T_H_2 cell-dependent lung M2 differentiation drives host resistance during primary infection with the parasitic nematode *Litomosoides sigmodontis*^52^. Unexpectedly, we found that depleting CD4 cell compartment using the anti-CD4mAb (GK1.5 clone) administered only after the initial infection had no impact on pathogen clearance, downmodulation of IL-17 inflammation or tissue repair, which was an initial indication that ILC2s had a vital role in anamnestic immunity

Bouchery and colleagues reported that lung T_H_2 cells and ILC2s work together to eliminate *Nb* lung larvae within 48h post re-challenge^25^. Similarly, our data propose a mechanism wherein IL-5/13+ ILC2s in the lung can aid T_H_2 cells in recruiting eosinophils and Arg1+ M2 macrophages for parasite elimination. Lung eosinophils bind to *Nb* larvae through complement and release their intracellular granules that damage larvae^53,54^. ILC2-deficient (*Lcr1*^-/-^) mice had high numbers of lung larvae, low IL-5, eosinophil and M2 macrophage levels after reinfection compared to controls. However, restoration of worm killing in *Lcr1*^-/-^ mice through Areg supplementation was marked by increased T_H_2 cell numbers and basophils, not M2 macrophages.

Our data corroborate extensive crosstalk between ILC2s and CD4^+^ T cells and the emerging “trained ILC2” concept. Continuous depletion of CD4^+^ T cells across primary and secondary infection significantly reduced ILC2 expansion and Type 2 cytokine production, but CD4 depletion only prior to secondary infection had little impact. It is likely that during initial antigen encounter, CD4+ T cells prime lung ILC2s for optimal efficacy against *Nb* reinfection. CD4+ T cell-derived IL-2 can promote ILC2 proliferation and Type 2 cytokine expression during primary *Nb* infection^29^ and IL-2 supplementation of CD4+ T cell-depleted mice restores parasite control and increases lung ILC2 abundance^25^. Therefore, the upregulation of IL-2rα transcript expressed by ILC2s after reinfection in our study (Fig. 3F) could indicate that CD4+ T cells through IL-2 secretion train lung ILC2s through reshaping their metabolic, transcriptional and/or epigenetic profiles to better respond to *Nb* rechallenge. Indeed, in an allergic asthma model CD25+ ILC2s exhibited long-term survival and enhanced cytokine secretion after secondary sensitization with the same allergen^22^. While this study did not delve into ILC2 subsets, reports that utilized Il1rl1-(IL-33 receptor) and Il17rb-(IL-25 receptor) deficient mice revealed that IL-33 and IL-25 were distinct in their ability to promote Type-2 cytokine responses or helminth clearance^55,56^. Ricardo-Gonzalez and colleagues extended this concept and distinguished two waves of ILC2s entering the bloodstream during *Nb* infection. At d5 post-infection, ILC2s are predominantly intestinal-derived IL-25R+ cells resembling inflammatory ILC2s (iILC2s)^18^ whereas at d12 p.i., ILC2s have a lung-derived phenotype expressing IL-33R and referred to as natural ILC2s (nILC2s)^57^. Small intestine iILC2s can migrate into lymph nodes to interact with primed CD4^+^T cells in an MHC class II-dependent manner^8,29^. Also, ILC2s provide help to T_H_2 cells via a PD-L1–PD-1-mediated interactions and the deletion of PD-L1 in ILC2s diminishes the capacity of T cells to produce type 2 cytokines or drive intestinal worm expulsion^7^. *Lcr1*^-/-^mice show impaired T_H_2 polarization following house dust mite antigen challenge ^31^ and we show that Areg can rescue this defect, consistent with ILC2-mediated T_H_2 cell expansion. This mouse strain was created through deletion of *Gata3* and *Rorα* binding sites essential for ILC2 development and expansion, but our data suggest the effects on T_H_2 cell in these mice are cell extrinsic.

Primary *Nb* infection causes severe, but transient lung hemorrhage due to migratory larvae that cross the pulmonary vasculature into alveolar space and this injury is further abbreviated upon re-challenge^45,46^. Lung hemorrhage is associated with IL-17A producing γδT cells and neutrophilia driven by Ym1+ macrophages, but Ym1 neutralization does not block worm clearance^11,12,47,58,59^. IL-4Rα signaling downmodulates IL-17 inflammation to allow emergence of Type 2 responses, restoration of pulmonary function via TFF2 and other IL-13-dependent reparative functions^60^. ILC2 functions are critical for suppressing excess pulmonary inflammation^61^ and we postulated that lack of ILC2-derived of Areg explained why continuous GK1.4 administration or gene-targeted deletion of ILC2 exacerbated IL-17+ γδT cell and neutrophil responses. Indeed, Areg can directly promote epithelial repair and myofibroblast differentiation to drive tissue repair in a variety of pathological contexts including dextran sodium sulfate-induced colitis and influenza virus infection^39,62^. Thus, it was surprising that exogenous rAreg administration did not reduce lung hemorrhage or block IL-17 associated inflammation.

Thus, we turned focus to Tryptophan Hydroxylase 1 (*Tph1*) which was moderately increased in trained lung ILC2s isolated after secondary infection. Tph1 is a rate-limiting enzyme for 5-HT production in peripheral tissues^63^ and intestinal Enterochromaffin cells are a major 5-HT source of this neurotransmitter^64^. Artis and colleagues demonstrated IL-33 is required for *Tph1* expression in ILC2 for adult worm expulsion from the intestine during primary *Nb* infection ^65^. Our data show that increased 5-HT levels in *Nb*-infected lung tissues were abrogated by all approaches where ILC2 were absent and they produced 5-HT in an IL-33 dependent manner. Strikingly, intranasal 5-HT administration blocked lung injury in an ILC2 and T_H_2 cell independent manner and Tph1 inhibition exacerbated the hemorrhage/IL-17 axis despite parasite clearance. 5-HT inhibition did not change lung ILC2 numbers (Fig. 5G) although they express the 5HT receptor *Htr1b*^65^. That 5-HT promotes wound healing in the skin^66–69^ opens a myriad of potential endothelial, epithelial, mesenchymal or hematopoietic cellular targets for serotonin activity explaining the immunoregulatory and reparative effects of 5-HT in damaged lung tissue.

In summary, this work supports the concept of trained ILC2, something that requires CD4+ T cell interactions during initial priming for optimal function during anamnestic Type 2 immunity. Trained ILC2 serve as a major source of 5-HT in hookworm infected lungs that limits tissue injury and prevents dysregulated inflammation. Investigation of whether 5-HT drives pulmonary tissue repair in other contexts is warranted.

## Materials and Methods

### Mice

Experimental groups ranged from 3-5 mice/group (matched for age and sex, with both sexes used), repeated at least twice to assure reproducibility and mice were tattooed for identification. All mice were 8-12 weeks old at the time of the experimentation. *Lcr1*^-/-^ (ILC2s-deficient) and Red5 mice (6(C)-Il5tm1.1(icre)Lky/J; AX strain #030926) mice were kindly donated by Henao-Mejia’s laboratory at University of Pennsylvania ^31^ and Paula Oliver at Children Hospital of Philadelphia respectively. CD45.1 C57bl6J mice (B6.SJL-Ptprca Pepcb/BoyJ; strain #002014) were purchased from the Jackson Laboratories. Breeding stock of wild-type C57Bl/6 (WT) mice were purchased from Taconic Laboratories. These mouse strains were bred in our animal facility at University of Pennsylvania School of Veterinary Medicine. All procedures were approved by the Institutional Animal Care and Use Committee of the University of Pennsylvania (protocol 805911). At the end of each experiment mice were euthanized by CO_2_ for all tissue recovery procedures following AVMA guidelines.

### CD4 antibody neutralization

For CD4+T cell depletion, mice were treated intraperitoneally (i.p.) with 1 mg or 0.5 mg of CD4 neutralizing antibody (GK.15, BioXcell, cat# BE0003-1) while control mice were treated with the same doses of Isotype antibody (LTF-2, BioXcell, cat#BE0090). In the continuous CD4+T cells depletion model, neutralizing antibody were given prior both primary (day -1) and secondary (day14) infections whereas for the delayed depletion model, mAb was administrated just prior the secondary infection (day 14 and day 21 p.i.).

### Amphiregulin and serotonin supplementation

For Areg supplementation, *Lcr1*^-/-^ mice were treated intranasally (i.n.) with 5µg of recombinant mouse amphiregulin protein (R&D Systems, cat# 989-AR-100) a day prior and two consecutive days following the secondary challenge with *N.b.* While an intranasal I.n. delivery of serotonin (5-HT) (Tocris Bioscience, cat# 3547) in 50 ul at 0.1 mg/mL was administered, accordingly 300 mg/kg of p-Chlorophenylalanine (Tocris Bioscience, cat# 0938) was injected i.p. to WT mice from day 14-day 19 post-primary infection

### Nippostrongylus brasiliensis Infection

Following sedation with up to 4 % isoflurane, mice were infected by subcutaneous injection of 650 *N. brasiliensis* iL3, performed twice in all experiments. Fecal egg burden was assessed using previously published methods ^70^. To ensure parasite clearance, the anthelminthic drug pyrantel pamoate was administered by oral gavage at the of 0.680 mg in 150µL/mouse after primary infection as indicated by “Rx” in experimental design figures to control parasite load across experimental groups.

### Adoptive transfer of CD4+ T cells

Single cell suspensions from the spleen, the mesenteric and the cervical lymph nodes were isolated from naïve CD45.1 mice, submitted to an initial enrichment using Naïve CD4+ T cells isolation kit (Miltenyi Biotech, Cat# 130-104-453) according to the manufacturer’s protocol. Enriched cell suspensions were subjected to a surface marker staining and viability determined. Live naive CD4+T cells (CD45^+^CD3^+^CD4^+^CD62L^+^) were sorted using the BD FACSARIA III cell sorter (BD Biosciences) and approximately 3 million cells were injected i.v. to *Lcr1*^-/-^ mice a day prior to primary *Nb* infection.

| Antibodies used for FACS sorting |  |  |
| --- | --- | --- |
| Fluorochrome/Markers (Dilution) | Manufacturer | #Catalog |
| FITC anti-mouse CD3 Antibody (1:100) | Biolegend | 100204 |
| PerCP/Cy5.5 anti-mouse CD4 Antibody (1:100) | Biolegend | 100434 |
| APC anti-mouse CD45 Antibody (1:100) | Biolegend | 147708 |
| PE/Dazzle 594 anti-mouse CD62L Antibody (1:100) | Biolegend | 104448 |
| DAPI (1: 10 000) | ThermoFisher | 62248 |

### Lung digestion and flow cytometry

Lungs were surgically dissected, dissociated and digested in a solution containing 0.15 mg/mL of Liberase (Sigma-Aldrich), 0.4 mg/mL of Dispase II (Sigma-Aldrich), and 0.014 mg/mL DNase I (Roche) dissolved in DMEM-containing 10% FBS for 40 mins at 37°C with constant agitation. Cell preparations were filtered twice through 100μm and 40μm filters and resuspended in RPMI supplemented with 10% FBS and incubated with Cell Activation Cocktail (with Brefeldin A) for 5-6hours. Lung cells were stained for live/dead cell exclusion using LIVE/DEAD™ Fixable Aqua Dead Cell Stain Kit following manufacturer’s protocol. Fc Block was performed for 25 mins at 4°C followed by surface marker staining for 25 mins on ice. Next, eBioscience™ Foxp3/Transcription Factor Staining Buffer Set was used according to manufacturer’s protocol. Intracellular staining was done for overnight at 4°C. Cells were analyzed with a BD Symphony A3 Lite (BD bioscience, USA). Fluorescence minus one (FMO) staining was used to establish reliable and reproducible gates for each marker.

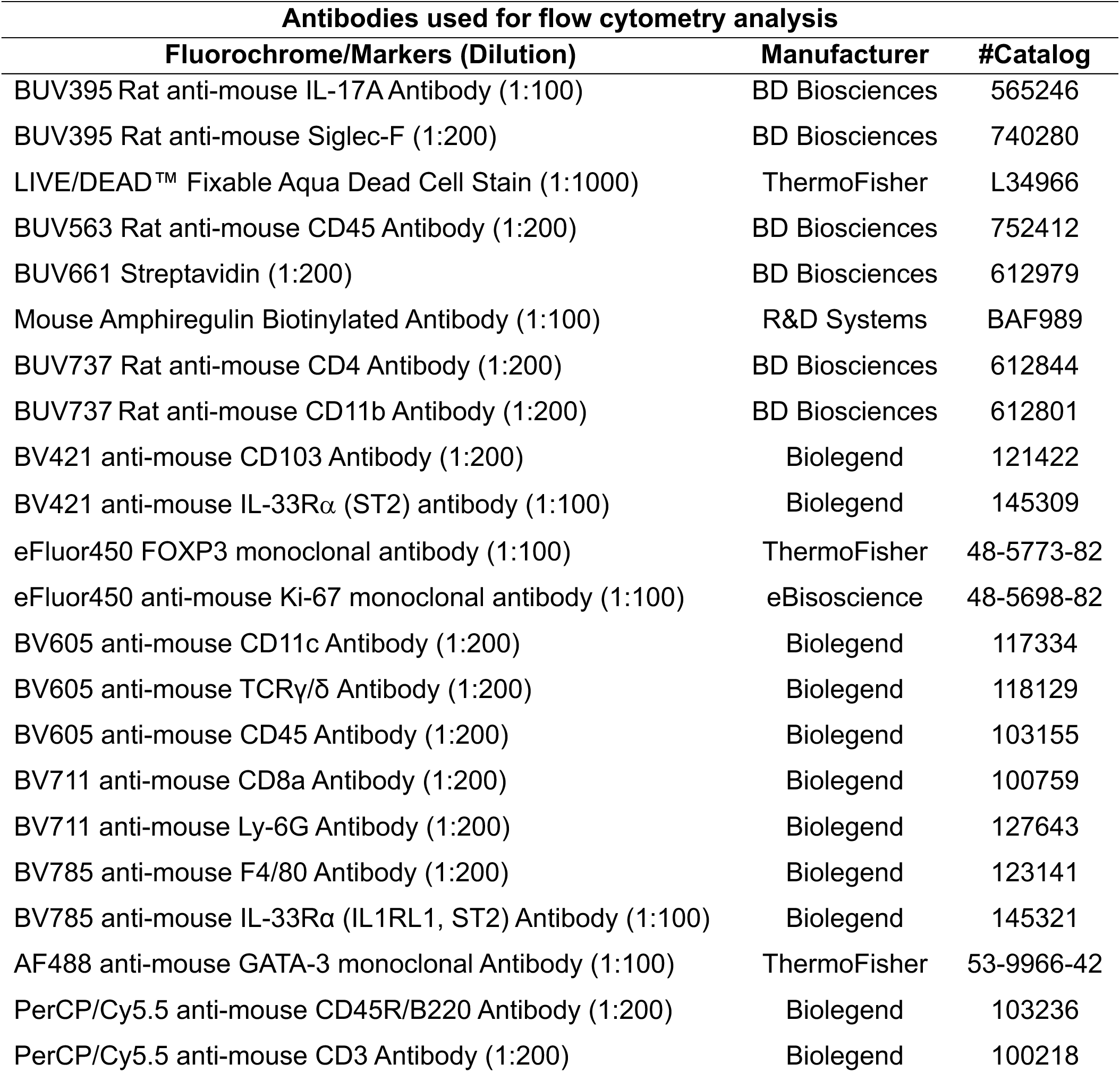
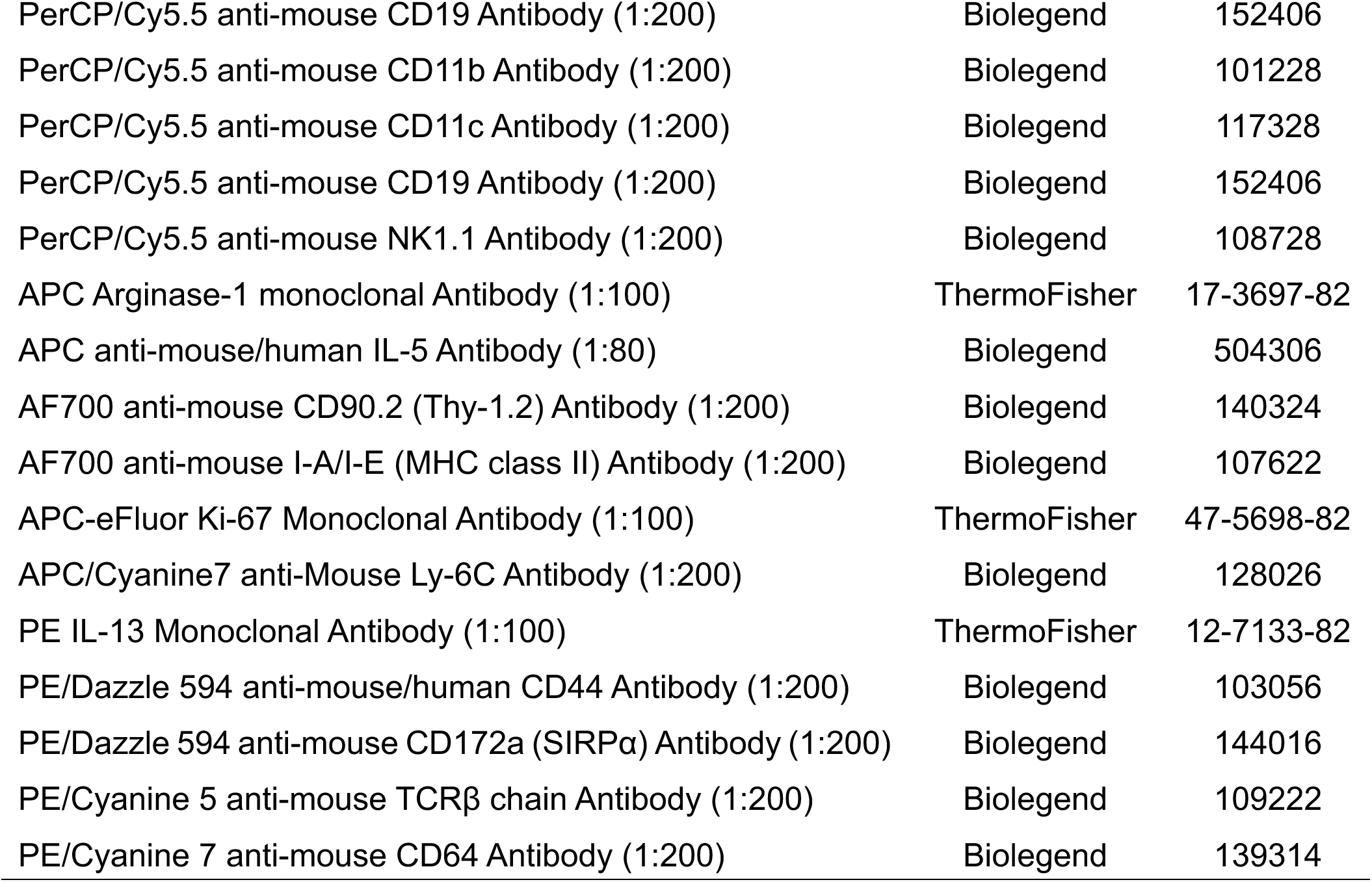

| **Antibodies used for flow cytometry analysis** |  |  |
| --- | --- | --- |
| **Fluorochrome/Markers (Dilution)** | **Manufacturer** | **#Catalog** |
| BUV395 Rat anti-mouse IL-17A Antibody (1:100) | BD Biosciences | 565246 |
| BUV395 Rat anti-mouse Siglec-F (1:200) | BD Biosciences | 740280 |
| LIVE/DEAD™ Fixable Aqua Dead Cell Stain (1:1000) | ThermoFisher | L34966 |
| BUV563 Rat anti-mouse CD45 Antibody (1:200) | BD Biosciences | 752412 |
| BUV661 Streptavidin (1:200) | BD Biosciences | 612979 |
| Mouse Amphiregulin Biotinylated Antibody (1:100) | R&D Systems | BAF989 |
| BUV737 Rat anti-mouse CD4 Antibody (1:200) | BD Biosciences | 612844 |
| BUV737 Rat anti-mouse CD11b Antibody (1:200) | BD Biosciences | 612801 |
| BV421 anti-mouse CD103 Antibody (1:200) | Biolegend | 121422 |
| BV421 anti-mouse IL-33Rα (ST2) antibody (1:100) | Biolegend | 145309 |
| eFluor450 FOXP3 monoclonal antibody (1:100) | ThermoFisher | 48-5773-82 |
| eFluor450 anti-mouse Ki-67 monoclonal antibody (1:100) | eBioscience | 48-5698-82 |
| BV605 anti-mouse CD11c Antibody (1:200) | Biolegend | 117334 |
| BV605 anti-mouse TCRγ/$\delta$ Antibody (1:200) | Biolegend | 118129 |
| BV605 anti-mouse CD45 Antibody (1:200) | Biolegend | 103155 |
| BV711 anti-mouse CD8a Antibody (1:200) | Biolegend | 100759 |
| BV711 anti-mouse Ly-6G Antibody (1:200) | Biolegend | 127643 |
| BV785 anti-mouse F4/80 Antibody (1:200) | Biolegend | 123141 |
| BV785 anti-mouse IL-33Rα (IL1RL1, ST2) Antibody (1:100) | Biolegend | 145321 |
| AF488 anti-mouse GATA-3 monoclonal Antibody (1:100) | ThermoFisher | 53-9966-42 |
| PerCP/Cy5.5 anti-mouse CD45R/B220 Antibody (1:200) | Biolegend | 103236 |
| PerCP/Cy5.5 anti-mouse CD3 Antibody (1:200) | Biolegend | 100218 |
| PerCP/Cy5.5 anti-mouse CD19 Antibody (1:200) | Biolegend | 152406 |
| PerCP/Cy5.5 anti-mouse CD11b Antibody (1:200) | Biolegend | 101228 |
| PerCP/Cy5.5 anti-mouse CD11c Antibody (1:200) | Biolegend | 117328 |
| PerCP/Cy5.5 anti-mouse CD19 Antibody (1:200) | Biolegend | 152406 |
| PerCP/Cy5.5 anti-mouse NK1.1 Antibody (1:200) | Biolegend | 108728 |
| APC Arginase-1 monoclonal Antibody (1:100) | ThermoFisher | 17-3697-82 |
| APC anti-mouse/human IL-5 Antibody (1:80) | Biolegend | 504306 |
| AF700 anti-mouse CD90.2 (Thy-1.2) Antibody (1:200) | Biolegend | 140324 |
| AF700 anti-mouse I-A/I-E (MHC class II) Antibody (1:200) | Biolegend | 107622 |
| APC-eFluor Ki-67 Monoclonal Antibody (1:100) | ThermoFisher | 47-5698-82 |
| APC/Cyanine7 anti-Mouse Ly-6C Antibody (1:200) | Biolegend | 128026 |
| PE IL-13 Monoclonal Antibody (1:100) | ThermoFisher | 12-7133-82 |
| PE/Dazzle 594 anti-mouse/human CD44 Antibody (1:200) | Biolegend | 103056 |
| PE/Dazzle 594 anti-mouse CD172a (SIRPα) Antibody (1:200) | Biolegend | 144016 |
| PE/Cyanine 5 anti-mouse TCRβ chain Antibody (1:200) | Biolegend | 109222 |
| PE/Cyanine 7 anti-mouse CD64 Antibody (1:200) | Biolegend | 139314 |

### Quantification of *Nippostrongylus* parasites in lung and intestine

For lung larvae quantification, after dissection lung were placed a 60 mm x 15 mm petri dish and minced using scissors into ∼1-2 mm pieces. Thereafter, 5-7 mL 1X PBS was immediately and incubated at 37°C for 2 hours. For adult worms, small intestinal tissue was longitudinally open and placed on a Baermann apparatus using metal sieves incubated for 4 hours at 37°C. *N. brasiliensis* L4 larvae or adult worms were counted under the light stereoscope.

### ELISA

Bronchoalveolar lavage fluid was spun down at 1500 rpm at 4°C for 5 minutes to isolate supernatant from cellular debris. BALF supernatants were used to measure cytokine concentration levels using the commercially kits listed in the following table, following the manufacturer’s instructions. A Biotech Synergy 2 Plate reader was used to determine absorbance, and concentrations were determined from the standard curve using a 4-parameter fit, which produced R^2^ values of greater than at least 0.9.

| Kit names | Manufacturers | #Catalog |
| --- | --- | --- |
| Mouse Amphiregulin | R&D systems | DYY989 |
| Mouse IL-5 | Fisher scientific | 88-7054-86 |
| Mouse IL-13 | Fisher scientific | 88-7137-88 |
| Serotonin Competitive ELISA | Fisher Scientific | EEL006 |

### Single-cell RNA sequencing

Lung ILC2s from WT mice were harvested and digested a two timepoints: day 3 post primary and day 3 post-secondary challenge with *N.b*. Cell suspensions were submitted to preliminary MACS enrichment following by FACS sorting and sort-purified ILC2 were identified as Live CD45^+^Lin^-^CD90.2^+^CD127^+^ using an BD FACS Aria II sorter (BD Biosciences). Sort-purified cells were resuspended in PBS supplemented with 10%FBS to achieve a target cell concentration of 700 to 1200 cells per μL. Cell viability was determined by trypan blue exclusion and only samples with >85% viability were processed. Next-generation sequencing libraries were prepared using the 10x Genomics Chromium Single Cell 3’ Reagent kit v3 (10X Genomics) per manufacturer’s instructions. Libraries were uniquely indexed using the Chromium dual Index Kit, pooled, and sequenced on an Illumina NovaSeq 6000 sequencer in a paired-end, dual indexing run. Sequencing for each library targeted 20,000 mean reads per cell. Data was then processed using the Cell Ranger pipeline (10x Genomics, v.6.1.2) for demultiplexing and alignment of sequencing reads to the mm10 transcriptome and creation of feature-barcode matrices. Data were further processed with Seurat 4.0 R package ^71^. For quality control, only genes expressed in at least 3 cells and cells expressing at least 200 genes were included. Cells expressing >10% mitochondrial genes were excluded from the downstream analysis. Data were normalized and scale using default parameters, and the number of principal components were estimated using *RunPCA* followed by *ElbowPlot*. Uniform Manifold Approximation and Projection (UMAP) was used for dimensionality reduction and performed using *RunUMAP*. Markers for cell clusters were identified using the *FindAllMarkers* function equipped in the Seurat and cell types were annotated manually using canonical markers. We deployed the processed results in Shiny App (https://reedlab3.shinyapps.io/HerbertD_NicolisI_ScV3_NEW/) for further exploration. The statistics were computed using the Wilcoxon rank-sum test in combination with limma model. The significant threshold was set up as avg_log2FC >0.5 or <-0.5 and the percentage of cells in each group >5% and FDR<=0.05

| **Antibodies used for MACS-enrichment and FACS-sorting** |  |  |
| --- | --- | --- |
| **Fluorochrome/Markers (Dilution)** | **Manufacturer** | **#Catalog** |
| Biotin anti-mouse CD3 Antibody (1:100) | Biolegend | 100244 |
| Biotin anti-mouse CD19 Antibody (1:200) | Biolegend | 115504 |
| Biotin anti-mouse/human CD11b Antibody (1:200) | Biolegend | 101204 |
| Biotin anti-mouse CD11c Antibody (1:100) | Biolegend | 117304 |
| Biotin anti-mouse CD31 Antibody (1:100) | Biolegend | 102504 |
| Biotin anti-mouse CD326 (EpCAM) Antibody (1:100) | ThermoFisher | 13-5791-82 |
| PerCP Cy5.5 anti-mouse CD3 (1:200) | Biolegend | 137609 |
| PerCP Cy5.5 anti-mouse CD19 (1:200) | Biolegend | 152406 |
| PerCP Cy5.5 anti-mouse CD11c (1:200) | Biolegend | 117328 |
| PerCP Cy5.5 anti-mouse CD11b (1:300) | Biolegend | 101228 |
| PerCP Cy5.5 anti-mouse NK1.1 (1:100) | Biolegend | 108728 |
| PerCP Cy5.5 anti-mouse TCRb (1:200) | Biolegend | 109228 |
| PerCP Cy5.5 anti-mouse CD5 (1:200) | Biolegend | 100624 |
| PerCP Cy5.5 anti-mouse Ter119 (1:200) | Biolegend | 116226 |
| APC anti-mouse CD45 Antibody (1:300) | Biolegend | 147708 |
| FITC anti-mouse CD90.2 Antibody (1:300) | Biolegend | 140303 |
| PE-Dazzle anti-mouse CD127 (1:100) | Biolegend | 135032 |
| DAPI Solution (1:10 000) | ThermoFisher | 62248 |

### ILC2 culture

FACS-sorted lung ILC2 from WT mice were obtained at three time points: naïve, day 3 post-primary *N. brasiliensis* infection, and day 3 post-secondary challenge. Sorted ILC2s were cultured in 96-well plates under standard conditions (5% CO₂, 37°C) for 7 days in ILC2 culture medium containing 50 ng/mL recombinant IL-2 (ThermoFisher, Cat# 212-12) and 50 ng/mL recombinant IL-7 (Biolegend, Cat# 577804) in RPMI-1640 supplemented with 10% FBS, 1% penicillin/streptomycin, and 55 μM 2-mercaptoethanol. Subsequently, 3 × 10⁵ cells were stimulated with 50 ng/mL recombinant IL-33 (Biolegend, Cat# 580504) for 48 hours and 5-HT levels in the culture supernatant quantified by ELISA.

### CD44+CD4+T cells sorting and culture

Lungs from WT mice (n = 10) subjected to *Nb* reinfection were enzymatically digested, and single-cell suspensions were enriched for CD4⁺ T cells using the CD4⁺ T Cell Isolation Kit (Miltenyi Biotec, Cat# 130-104-453) according to the manufacturer’s protocol. Enriched cells were stained with surface markers and viability dye and antigen-experienced CD4⁺ T cells (Live CD45⁺Lin⁻CD3⁺CD4⁺CD44⁺ cells) were sorted on a BD FACSAria Fusion cell sorter (BD Biosciences). Sorted CD44⁺CD4⁺ T cells were cultured in anti-CD3ε–coated 96-well plates (2 μg/mL in PBS; Invitrogen, Cat# 16-0031-82) in TH2-polarizing medium containing 0.5 μg/mL anti-CD28 (Biolegend, Cat# 102116), 1 μg/mL anti–IFN-γ (Biolegend, Cat# 505834), 50 ng/mL rIL-2 (ThermoFisher, Cat# 212-12), and 50 ng/mL rIL-4 (ThermoFisher, Cat# 212-12) in RPMI-1640 supplemented with 10% FBS, 1% penicillin–streptomycin, 1:100 non-essential amino acids, and 55 μM 2-mercaptoethanol. After 7 days of culture (37°C, 5% CO₂), 3 × 10⁵ cells were stimulated with r IL-33 (50 ng/mL) or amphiregulin (rAreg, 100 ng/mL) for 48 h, followed by serotonin quantification in supernatants by ELISA or intracellular analysis by flow cytometry.

| Antibodies used for FACS sorting |  |  |
| --- | --- | --- |
| Fluorochrome/Markers (Dilution) | Manufacturer | #Catalog |
| FITC anti-mouse CD3 Antibody (1:100) | Biolegend | 100204 |
| BV421 CD4 Antibody (1:100) | Biolegend | 100434 |
| PE anti-mouse CD45 Antibody (1:100) | Biolegend | 147708 |
| PE/Dazzle 594 anti-mouse CD44 Antibody (1:100) | Biolegend | 104448 |
| PerCP Cy5.5 anti-mouse CD19 (1:200) | Biolegend | 152406 |
| PerCP Cy5.5 anti-mouse CD11c (1:200) | Biolegend | 117328 |
| PerCP Cy5.5 anti-mouse CD11b (1:200) | Biolegend | 101228 |
| PerCP Cy5.5 anti-mouse Ter119 (1:200) | Biolegend | 116226 |
| PerCP Cy5.5 anti-mouse NK1.1 (1:200) | Biolegend | 108728 |
| Live/Dead™ Fixable Near-IR Dead Cell Stain | Invitrogen/Thermofisher | L34975 |

### Hematoxylin and eosin staining

At day 3 postsecondary challenge with *Nb*, lung tissues were collected from mice infection, immediately perfused and fixed in 4%Paraformaldehyde for 48 hours 4°C. Fixed tissues were then dehydrated through a graded series of ethanol, cleared in xylene, and embedded in paraffin. Serial 5-µm-thick sections were cut using a microtome and mounted on glass slides. For Hematoxylin and Eosin (H&E) staining, sections were deparaffinized in xylene, rehydrated through descending grades of ethanol to distilled water, and stained with Harris hematoxylin for 5 minutes to visualize nuclei. Slides were rinsed in running tap water, differentiated briefly in 1% acid alcohol, and counterstained with 1% eosin Y solution for 2 minutes to visualize cytoplasmic and extracellular components. After dehydration and clearing, slides were mounted with DPX mounting medium and examined under a bright-field microscope (Nikon Eclipse Ti2).

### Immunostaining

Immunostaining on the lung was performed as previously described ^72^. In detail, After Red5 mice were euthanized, lungs perfused with 10 to 15 mL cold PBS through the right ventricle, followed by fixation by instillation with 4 mL 4% paraformaldehyde (PFA) through the trachea. The collected tissue was further fixed in PFA for 1h at 4°C, washed in PBS three times, dehydrated in 30% sucrose overnight at 4°C, and embedded in OCT (Sakura) for cryo-sectioning. 15 µm-thick cryo-sections were washed in PBS, incubated in blocking buffer with 5% normal donkey serum (Jackson Immunoresearch) and 0.1% Triton X-100 in PBS for 30 minutes at RT, followed by overnight primary Ab incubation, secondary antibody incubation was performed for 2h and DAPI for 15 min. Each incubation was followed by a 3X washing step with a solution of 1% BSA in 1XTBS for 5 minutes. Images were captured using Leica DMI 6000 inverted confocal microscope (Leica Microsystems, USA). Each cell type was quantified on fifteen non-overlapping high-power fields per section mouse.

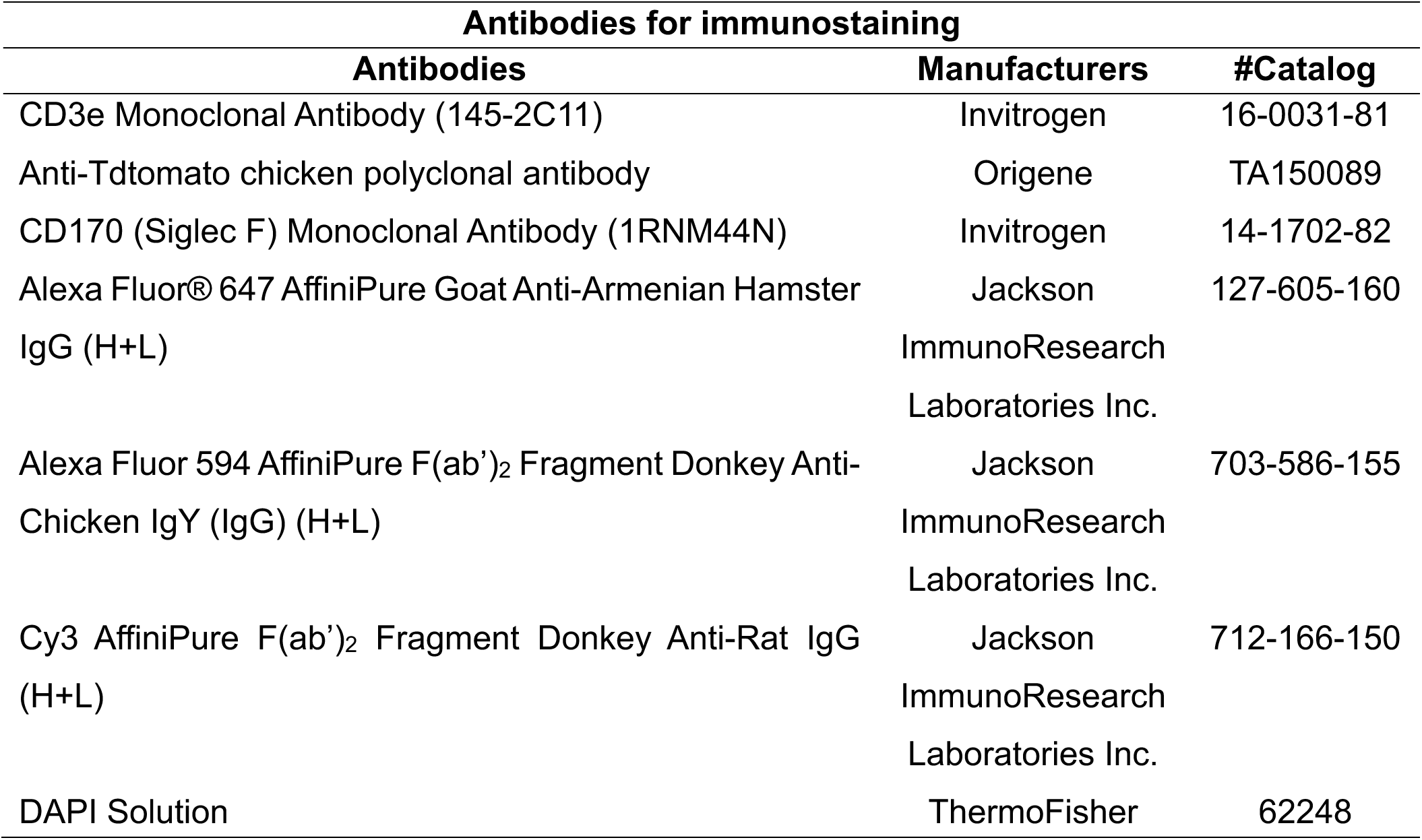

### Statistics

All data are displayed as mean ± SEM. P < 0.05 was considered significantly different. Because these data were normally distributed, parametric statistical tests were used. Statistical analysis was performed using Student’s t-test for two groups or one- or two-way ANOVA for three groups with Tukey multiple comparison test as post-hoc tests in Prism 10 (GraphPad Software, USA). For scRNA-Seq analysis, the statistics were computed using the Wilcoxon rank-sum test in combination with limma model. The significant threshold was set up as avg_log2FC >0.5 or <-0.5 and the percentage of cells in each group >5% and FDR<=0.05.

## Supporting information

Graphical Abstract

## Data availability

All data needed to draw conclusions are presented in the article and supplementary materials. Transcriptome data are available online at GEO repository GSE295590.

## Credit authorship contribution statement

**Ulrich M. Femoe**: Conceptualization, Investigation, Methodology, Data curation, Visualization, Writing the original draft, Editing; **Fungai Musaigwa**: Investigation, Methodology, Review and Editing; **Imani Nicolis**: Investigation, Methodology; **Li-Yin Hung**: Investigation, Methodology and Review and Editing; **Chinwekele Uzoije**: Investigation, Methodology; **Camila Napuri**: Investigation, Methodology; **Heather L. Rossi**: Data curation, Project administration, Funding acquisition, Review and editing; **Cailu Lin**: Data curation, Visualization, Investigation, Methodology; **Heidi Winters**: Investigation, Methodology; **Danielle R. Reed**: Data curation, Investigation, Methodology; **Juan M. Inclan-Rico**: Data curation, Funding acquisition Supervision, Review and editing; **De’Broski R. Herbert**: Conceptualization, Data curation, Project administration, Supervision, Funding acquisition, Review and editing the final draft.

## Acknowledgements

We thank the Henao-mejia and Oliver laboratories, for providing the *Lcr1*^-/-^ and Red5 mouse strains. We thank the University of Pennsylvania CDB Microscopy Core (RRID SCR_022373) for providing training on confocal imaging.

---

**Supplemental Figure 1. Impact of different CD4+T cell depletion strategies on primary infection and flow cytometry gating strategy. A)** *Nb* fecal egg counts of WT mice during primary infection subjected to the continuous CD4 T cell depletion strategy or **(B)** the delayed CD4 depletion strategy. Rx indicates oral gavage with pyrantel pamoate **(C)** Flow cytometry gating strategy including fluorescence minus one (FMO) controls for identification of lung tissue lymphocyte populations at 3 days post-secondary infection. CD4+T cell population= Live CD45^+^Lin^-^ CD90.2^+^TCRβ^+^TCRγδ^-^CD8^-^CD4^+^; T_H_2 cell population: LiveCD4^+^FoxP3^-^CD44^+^GATA3^+^; Total ILC2 population: Live CD45^+^Lin^-^CD90.2^+^TCRβ^-^TCRγδ^-^GATA3^+^ cells; IL-5+IL-13+ILC2 population: ILC2^+^IL-5^+^IL-13^+^; γδT cell population: Live CD45^+^Lin^-^CD90.2^+^ TCRβ^-^ TCRγδ^+^; IL-17A+γδT cell population: γδT cells^+^IL-17A^+^ cells. **(D)** Gating strategy for lung myeloid cell populations at 3 days post-secondary infection. Eosinophil population: Live CD45^+^Lin^-^CD11b^+^Ly6C^-^Ly6G^-^CD64^+^Siglec-F^+^; Arginase-1+macrophage(M2) population: Live CD45^+^Lin^-^CD11b^+^Ly6C^-^Ly6G^-^Siglec-F^-^CD64^+^Arg1^+^; and Neutrophil population: Live CD45^+^Lin^-^CD11b^+^Ly6C^+^Ly6G^+^.

**Supplemental Figure 2. Impact of continuous vs. delayed CD4+T cell depletion strategy on lung CD4, IL-13 levels, ILC2, γδT cell, M2 macrophage, eosinophil, and neutrophil populations following *Nb* re-infection. (A)** Representative flow cytometry contour plots showing numbers of lung tissue CD4+T cells at 3 days post-secondary infection during continuous vs. **(B)** delayed treatment with anti-CD4 mAb (clone GK1.5). **(C)** IL-13 levels in BAL fluid as determined by ELISA during the continuous vs. **(D)** delayed CD4 depletion strategy. **(E)** Representative contour plots and frequency of lung eosinophils in continuous vs. **(F)** delayed CD4 depletion strategy. **(G)** Representative contour plots and total number of IL-5+IL-13+ lung ILC2 in continuous vs. **(H)** delayed CD4 depletion strategy. **(I)** Representative contour plots showing frequency of Arginase-1+ lung tissue macrophages in continuous vs. **(J)** delayed CD4 depletion strategy. **(K)**, Representative contour plots showing numbers of total lung γδT cells during continuous vs. **(L)** delayed CD4 depletion strategy. **(M)** Representative contour plots showing frequency of IL-17A producing γδT cells during continuous vs. **(N)** delayed CD4 depletion strategy. **(O)** Representative contour plots showing frequency of lung tissue neutrophils during continuous vs. **(P)** delayed CD4 depletion strategy. Mean and standard error shown. *P* values were determined by two-tailed Student’s t-test. *P<0.05, **P<0.01, ***P<0.001 ****P<0.0001, ns: non-statistically significant. Representative of two independent experiments.

**Supplemental Figure 3. *Lcr1* deficient mice have basal increase in lung γδ T cells and adoptive transfer of CD4+T cells increases basophils but fails to promote Type 2 responses or reduce IL-17 producing γδ T cell responses upon *Nb* re-infection. (A)** Representative flow cytometry contour plots showing the frequency and total cell number of the lung γδT cell population in non-infected WT vs *Lcr1*^-/-^ naïve mice. **(B)** Fecal egg counts during primary *Nb* infection of WT vs. *Lcr1*^-/-^ mice. **(C)** Fecal egg counts during primary *Nb* infection comparing WT vs. *Lcr1*^-/-^ vs. *Lcr1*^-/-^ mice given an adoptive transfer of 3 x 10^6^ naïve CD4+T cells on day -1. Rx indicates oral gavage with pyrantel pamoate. **(D)** IL-5 levels in BAL fluid from mice described in “C” analyzed at 3 days post-secondary infection. **(E)** IL-13 levels in BAL fluid from mice described in “C” analyzed at 3 days post-secondary infection. **(F)** Representative flow cytometry contour plots showing the frequency **(G)** lung tissue eosinophils, **(H)** lung basophils, **(I)** both frequency and total number of total γδ T cells **(J)** frequency of IL-17A producing γδT cells and **(K)** frequency of lung neutrophils on day 3 post-secondary infection of mice described in “C”. Mean and standard error values shown. *P* values were determined by one-way ANOVA followed by Tukey post-hoc test. *P<0.05, **P<0.01, ***P<0.001, ns: non-statistically significant. Representative of two independent experiments.

**Supplemental Figure 4. Abundance of T_H_2 and ILC2 in lung tissues of naïve and *Nb*-infected IL-5 td-Tomato fluorescent reporter (Red5) mice. (A)** Schematic shows timepoints for lung immunostaining after primary vs. secondary *Nb* infection in the Red5 strain (n=3/group) **(B, C)** Representative fluorescent microscopy images for control (no primary Ab) or single Ab fluorescence immunostaining for DAPI, CD3, Siglec-F and Td-tomato with merged image shown for each. **(D)** Representative merged images for Red5 mice that were naïve or 3 days post-primary, and 3 days and post-secondary infection. Left column is 40X magnification and inset shown on right is 63x magnification. Scale bar=20 microns **(E)** Quantification of lung eosinophils, T_H_2 and ILC2 at timepoints indicated in “D” **(F)** Quantification of lung ILC2s vs. T_H_2 by intracellular at timepoints indicated in “D”. **(G-I)** Quantification of lung IL-5+, IL-13+ and IL-5+IL13+ ILC2 vs T_H_2 cells at 3 days post-secondary infection. For IFA staining, P values were determined by two-way ANOVA for grouped data set or One-way ANOVA for non-grouped data sets followed by Tukey post-hoc multiple comparison test. *P<0.05, **P<0.01, ***P<0.001, ****P<0.0001, ns: non-statistically significant

**Supplemental Figure 5. Areg supplementation increases T_H_2 cells, IL-13, eosinophils and M2 macrophages, but not does not block γδT cell responses. (A)** Representative flow cytometry contour plots show frequencies of T_H_2 cells in lung tissues of *Nb*-infected for WT, *Lcr1*^-/-^ and rAreg supplemented *Lcr1*^-/-^ mice. **(B)** Levels of IL-13 in BAL fluid as determined by ELISA. **(C)** Representative contour plots show numbers of lung eosinophils **(D)** numbers of lung Arginase-1+macrophages, E) numbers of total lung γδT cells and **(F)** frequency of lung IL-17 producing γδT cells. All analysis performed at 3 days post-secondary infection. Mean and standard error values shown. *P* values were determined by one-way ANOVA followed by Tukey post-hoc multiple comparison test. *P<0.05, **P<0.01, ****P<0.0001, ns: non-statistically significant. Representative of two independent experiments.

**Supplemental Figure 6. Areg drives proliferative expansion of T_H_2 cells (A)** Flow cytometry gating strategy to identify live CD45^+^TCRβ^+^CD4^+^ cells following MACS-enrichment and FACS sorting prior to culture with recombinant amphiregulin or vehicle. **(B-C)** t-SNE dimensionality reduction analysis for specified markers for identification of T_H_2 subsets following culture with recombinant amphiregulin or vehicle. **(D)** FlowSOM clustering overlaid on t-SNE dimensionality reduction and FlowSOM cluster sizes. **(E)** Representative flow cytometry contour plots show numbers of GATA3 expressing T_H_2 cels, **(F)** Ki-67^+^ T_H_2 cells, and **(G)** IL-5+ and IL-13+ T_H_2 cells after culture with rAreg or vehicle control. Mean and standard error shown. *P* values were determined by two-tailed Student’s t-tests. *P<0.05, **P<0.01 ns: non-statistically significant. Representative of two independent experiments.

**Supplemental Figure 7. Serotonin (5-HT) is produced by ILC2 and controls total γδT cell, IL-17+ γδ T cell and neutrophil responses in lung tissues of *Nb* re-infected mice (A)** Volcano plot shows increased Tph1 expression in trained lung nILC2s vs. primary lung nILC2 as determined by sc-RNAseq analysis. **(B)** Schematic for lung ILC2 isolation following MACS-enrichment, FACS-sort and 7-day culture. **C)** Comparison of lung ILC2s from naïve, primary infected, re-infected mice and memory T_H_2 cells (300,000 cells/well) in the presence or absence of recombinant IL-33 (rIL-33) for 48h. Levels of 5-HT in culture supernatant determined by ELISA. **(D)** Representative flow cytometry contour plots showing frequency of total lung ILC2 **(E)** total number of γδT cells, **(F)** frequency of IL-17+ γδT cells and **(G)** frequency of neutrophils. Data shown in **(D-G)** are from mice subjected to the continuous CD4 depletion strategy and supplemented with 5-HT or the Tph1 inhibitor PCPA. Mean and standard error shown. *P* values were determined by two-tailed Student’s t-test for the comparison between two groups or one-way ANOVA followed by Tukey post-hoc test for comparison involving more than two groups. *P<0.05, **P<0.01, ***P<0.001, ****P<0.0001, ns: non-statistically significant. Representative of two independent experiments.

## References

1. McDaniel, M. M., Lara, H. I. & Moltke, J. von Initiation of type 2 immunity at barrier surfaces. Mucosal Immunol. 16, 86–97 (2023).

2. Hung, L. Y. et al. IL-33 drives biphasic IL-13 production for noncanonical Type 2 immunity against hookworms. Proc. Natl. Acad. Sci. U. S. A. 110, 282–287 (2013).

3. Yasuda, K., Adachi, T., Koida, A. & Nakanishi, K. Nematode-infected mice acquire resistance to subsequent infection with unrelated nematode by inducing highly responsive group 2 innate lymphoid cells in the lung. Front. Immunol. 9, 1– 15 (2018).

4. Mi, L. L. & Guo, W. W. Crosstalk between ILC2s and Th2 CD4+T Cells in Lung Disease. J. Immunol. Res. 2022, (2022).

5. Gurram, R. K. & Zhu, J. Orchestration between ILC2s and Th2 cells in shaping type 2 immune responses. Cell. Mol. Immunol. 16, 225–235 (2019).

6. Qin, M. et al. Tissue microenvironment induces tissue specificity of ILC2. Cell Death Discov. 2024 101 10, 1–10 (2024).

7. Schwartz, C. et al. ILC2s regulate adaptive Th2 cell functions via PD-L1 checkpoint control. J. Exp. Med. 214, 2507–2521 (2017).

8. Mirchandani, A. S. et al. Type 2 Innate Lymphoid Cells Drive CD4+ Th2 Cell Responses. J. Immunol. 192, 2442–2448 (2014).

9. Schneider, C., et al. Tissue-Resident Group 2 Innate Lymphoid Cells Differentiate by Layered Ontogeny and In Situ Perinatal Priming. Immunity 50, 1425–1438.e5 (2019).

10. Paul, W. E. & Zhu, J. How are TH2-type immune responses initiated and amplified? Nat. Rev. Immunol. 10, 225–235 (2010).

11. Sutherland, T. E. et al. Chitinase-like proteins promote IL-17-mediated neutrophilia in a tradeoff between nematode killing and host damage. Nat. Immunol. 2014 1512 15, 1116–1125 (2014).

12. Chenery, A. L. et al. IL-13 deficiency exacerbates lung damage and impairs epithelial-derived type 2 molecules during nematode infection. Life Sci. Alliance 4, 1–14 (2021).

13. Thawer, S. G. et al. Lung-resident CD4+ T cells are sufficient for IL-4Rα-dependent recall immunity to Nippostrongylus brasiliensis infection. Mucosal Immunol. 2014 72 7, 239–248 (2013).

14. Herbert, D. R., Orekov, T., Perkins, C., Rothenberg, M. E. & Finkelman, F. D. IL-4Rα Expression by Bone Marrow-Derived Cells Is Necessary and Sufficient for Host Protection against Acute Schistosomiasis. J. Immunol. 180, 4948–4955 (2008).

15. Hung, L. Y. et al. Trefoil Factor 2 Promotes Type 2 Immunity and Lung Repair through Intrinsic Roles in Hematopoietic and Nonhematopoietic Cells. Am. J. Pathol. 188, 1161–1170 (2018).

16. Harvie, M., Camberis, M. & Gros, G. Le Development of CD4 T Cell Dependent Immunity Against N. brasiliensis Infection. Front. Immunol. 4, 74 (2013).

17. Harvie, M. et al. The Lung Is an Important Site for Priming CD4 T-Cell-Mediated Protective Immunity against Gastrointestinal Helminth Parasites. Infect. Immun. 78, 3753 (2010).

18. Huang, Y. & Paul, W. E. Inflammatory group 2 innate lymphoid cells. Int. Immunol. 28, 23–28 (2016).

19. Martinez-Gonzalez, I. et al. ILC2 memory: Recollection of previous activation. Immunol. Rev. 283, 41–53 (2018).

20. Wang, X., Peng, H. & Tian, Z. Innate lymphoid cell memory. Cell. Mol. Immunol. 2019 165 16, 423–429 (2019).

21. Martinez-Gonzalez, I. et al. Allergen-Experienced Group 2 Innate Lymphoid Cells Acquire Memory-like Properties and Enhance Allergic Lung Inflammation. Immunity 45, 198–208 (2016).

22. Jing, X. et al. The formation of memory-like innate lymphoid cells 2 in allergic asthma. J. Immunol. 198, 194.17–194.17 (2017).

23. Wilhelm, C. et al. Critical role of fatty acid metabolism in ILC2-mediated barrier protection during malnutrition and helminth infection. J. Exp. Med. 213, 1409– 1418 (2016).

24. Yu, H., Jacquelot, N. & Belz, G. T. Metabolic features of innate lymphoid cells. J. Exp. Med. 219, (2022).

25. Bouchery, T. et al. ILC2s and T cells cooperate to ensure maintenance of M2 macrophages for lung immunity against hookworms. Nat. Commun. 6, 1–13 (2015).

26. Hung, L.-Y. et al. Cell-Intrinsic Wnt4 Influences Conventional Dendritic Cell Fate Determination to Suppress Type 2 Immunity. J. Immunol. 203, 511–519 (2019).

27. Gause, W. C., Wynn, T. A. & Allen, J. E. Type 2 immunity and wound healing: evolutionary refinement of adaptive immunity by helminths. Nat. Rev. Immunol. 2013 138 13, 607–614 (2013).

28. Gurram, R. K. et al. Crosstalk between ILC2s and Th2 cells varies among mouse models. Cell Rep. 42, 112073 (2023).

29. Oliphant, C. J. et al. MHCII-Mediated Dialog between Group 2 Innate Lymphoid Cells and CD4+ T Cells Potentiates Type 2 Immunity and Promotes Parasitic Helminth Expulsion. Immunity 41, 283 (2014).

30. Klose, C. S. N. et al. The neuropeptide neuromedin U stimulates innate lymphoid cells and type 2 inflammation. Nature 549, 282–286 (2017).

31. Michieletto, M. F. et al. Multiscale 3D genome organization underlies ILC2 ontogenesis and allergic airway inflammation. Nat. Immunol. 24, 42–54 (2023).

32. Ohnmacht, C. & Voehringer, D. Basophils protect against reinfection with hookworms independently of mast cells and memory Th2 cells. J. Immunol. 184, 344–350 (2010).

33. Ohnmacht, C. et al. Basophils Orchestrate Chronic Allergic Dermatitis and Protective Immunity against Helminths. Immunity 33, 364–374 (2010).

34. Nussbaum, J. C. et al. Type 2 innate lymphoid cells control eosinophil homeostasis. Nature 502, 245–248 (2013).

35. Asaoka, M., Kabata, H. & Fukunaga, K. Heterogeneity of ILC2s in the Lungs. Front. Immunol. 13, 918458 (2022).

36. Halim, T. Y. F. Group 2 innate lymphoid cells in disease. Int. Immunol. 28, 13–22 (2016).

37. Thawer, S. G. et al. Lung-resident CD4+ T cells are sufficient for IL-4R-dependent recall immunity to Nippostrongylus brasiliensis infection. Mucosal Immunol. 7, 239–248 (2014).

38. Minutti, C. M. et al. A Macrophage-Pericyte Axis Directs Tissue Restoration via Amphiregulin-Induced Transforming Growth Factor Beta Activation. Immunity 50, 645–654.e6 (2019).

39. Monticelli, L. A. et al. Innate lymphoid cells promote lung-tissue homeostasis after infection with influenza virus. Nat. Immunol. 12, 1045–1054 (2011).

40. Singh, S. S. et al. Amphiregulin in cellular physiology, health, and disease: Potential use as a biomarker and therapeutic target. J. Cell. Physiol. 237, 1143– 1156 (2022).

41. Vergnes, M. Induction du comportement d’agression Rat-Souris par la p-chlorophenylalanine: Rôle de l’amygdale. Physiol. Behav. 25, 353–356 (1980).

42. Eberl, G., Colonna, M., Santo, J. P. D. & McKenzie, A. N. J. Innate lymphoid cells: A new paradigm in immunology. Science (80-.). 348, (2015).

43. Vivier, E. et al. Innate Lymphoid Cells: 10 Years On. Cell 174, 1054–1066 (2018).

44. Colonna, M. Innate Lymphoid Cells: Diversity, Plasticity and Unique Functions in Immunity. Immunity 48, 1104 (2018).

45. Craig, J. M. & Scott, A. L. Helminths in the lungs. Parasite Immunol. 36, 463–474 (2014).

46. Reece, J. J. et al. Hookworm-induced persistent changes to the immunological environment of the lung. Infect. Immun. 76, 3511–3524 (2008).

47. Weatherhead, J. E. et al. Host Immunity and Inflammation to Pulmonary Helminth Infections. Front. Immunol. 11, 594520 (2020).

48. Harvie, M., Camberis, M. & Gros, G. Le Development of CD4 T cell dependent immunity against N. brasiliensis infection. Front. Immunol. 4, 1–5 (2013).

49. Anthony, R. M. et al. Memory TH2 cells induce alternatively activated macrophages to mediate protection against nematode parasites. Nat. Med. 12, 955 (2006).

50. Chen, F. et al. Neutrophils prime a long-lived effector macrophage phenotype that mediates accelerated helminth expulsion. Nat. Immunol. 15, 938–946 (2014).

51. Chen, F. et al. Helminth resistance is mediated by differential activation of recruited monocyte-derived alveolar macrophages and arginine depletion. Cell Rep. 38, (2022).

52. Finlay, C. M. et al. T helper 2 cells control monocyte to tissue-resident macrophage differentiation during nematode infection of the pleural cavity. Immunity 56, 1064–1081.e10 (2023).

53. Giacomin, P. R. et al. The role of complement in innate, adaptive and eosinophil-dependent immunity to the nematode Nippostrongylus brasiliensis. Mol. Immunol. 45, 446–455 (2008).

54. Yasuda, K. & Kuroda, E. Role of eosinophils in protective immunity against secondary nematode infections. Immunol. Med. 42, 148–155 (2019).

55. Miller, M. M. & Reinhardt, R. L. The Heterogeneity, Origins, and Impact of Migratory iILC2 Cells in Anti-helminth Immunity. Front. Immunol. 11, 1–14 (2020).

56. Neill, D. R. et al. Nuocytes represent a new innate effector leukocyte that mediates type-2 immunity. Nature 464, 1367–1370 (2010).

57. Ricardo-Gonzalez, R. R. et al. Tissue-specific pathways extrude activated ILC2s to disseminate type 2 immunity. J. Exp. Med. 217, (2020).

58. Ajendra, J. et al. IL-17A both initiates, via IFNγ suppression, and limits the pulmonary type-2 immune response to nematode infection. Mucosal Immunol. 2020 136 13, 958–968 (2020).

59. Chen, F. et al. B Cells Produce the Tissue-Protective Protein RELMα during Helminth Infection, which Inhibits IL-17 Expression and Limits Emphysema. Cell Rep. 25, 2775–2783.e3 (2018).

60. Sutherland, T. E. et al. Ym1 induces RELMα and rescues IL-4Rα deficiency in lung repair during nematode infection. PLoS Pathog. 14, (2018).

61. Inclan-Rico, J. M. et al. Basophils prime group 2 innate lymphoid cells for neuropeptide-mediated inhibition. Nat. Immunol. 21, 1181–1193 (2020).

62. Monticelli, L. A. et al. IL-33 promotes an innate immune pathway of intestinal tissue protection dependent on amphiregulin-EGFR interactions. Proc. Natl. Acad. Sci. U. S. A. 112, 10762–10767 (2015).

63. Walther, D. J. & Bader, M. A unique central tryptophan hydroxylase isoform. Biochem. Pharmacol. 66, 1673–1680 (2003).

64. Mawe, G. M. & Hoffman, J. M. Serotonin Signaling in the Gastrointestinal Tract: Functions, dysfunctions, and therapeutic targets. Nat. Rev. Gastroenterol. Hepatol. 10, 473 (2013).

65. Flamar, A. L. et al. Interleukin-33 induces the enzyme tryptophan hydroxylase 1 to promote inflammatory group 2 innate lymphoid cell-mediated immunity. Immunity 52, 606 (2020).

66. Malinin, A., Oshrine, B. & Serebruany, V. Treatment with selective serotonin reuptake inhibitors for enhancing wound healing. Med. Hypotheses 63, 103–109 (2004).

67. Herr, N., Bode, C. & Duerschmied, D. The Effects of Serotonin in Immune Cells. Front. Cardiovasc. Med. 4, 1–11 (2017).

68. Gupta, D., Kaushik, D. & Mohan, V. Role of neurotransmitters in the regulation of cutaneous wound healing. Exp. Brain Res. 240, 1649–1659 (2022).

69. Sadiq, A. et al. 5-HT1A Receptor Function Makes Wound Healing a Happier Process. Front. Pharmacol. 9, 1–13 (2018).

70. Barreau, F. et al. Neonatal maternal deprivation promotes Nippostrongylus brasiliensis infection in adult rats. Brain. Behav. Immun. 20, 254–260 (2006).

71. Butler, A., Hoffman, P., Smibert, P., Papalexi, E. & Satija, R. Integrating single-cell transcriptomic data across different conditions, technologies, and species. Nat. Biotechnol. 36, 411–420 (2018).

72. Lv, Z., Liu, Z., Liu, Kuo Lin, Xiuyu Pu, Wenjuan Li, Yan Zhao, Huan Xi, Ying Sui, P., Vaughan, A. E. & Gillich, A. Alveolar regeneration by airway secretory-cell-derived p63+ progenitors. Stem Cell 31, 1–16 (2024).

