## Supplementary figures and images for "Trained ILC2 prevent IL-17-associated lung injury during helminth infection through a serotonin-dependent mechanism"

### Graphical Abstract

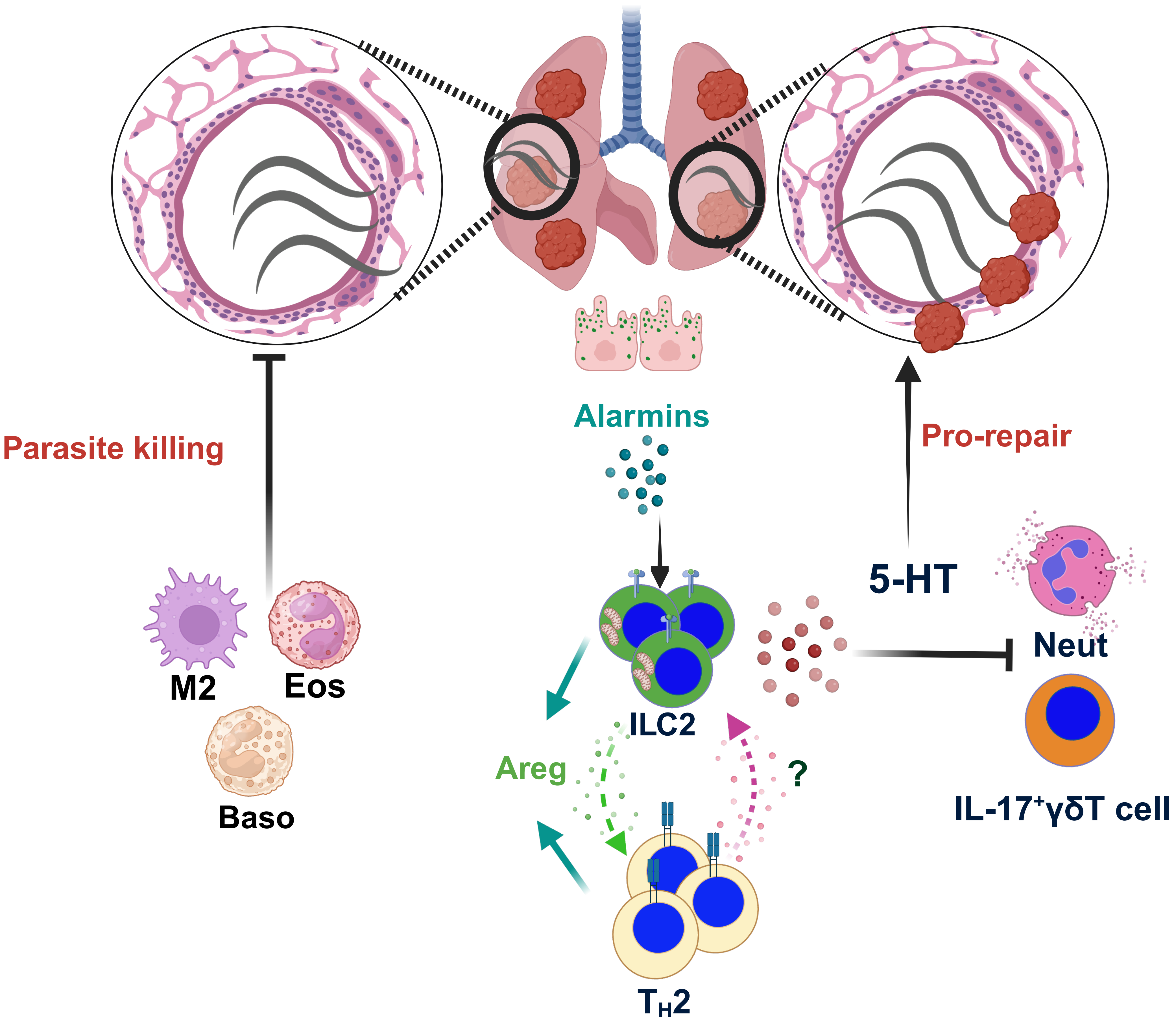


Created in BioRender. Femoe Membe, U. (2025) https://BioRender.com/ptwfqn9

### Supplemental Figure1

Fig. Supplemental 1

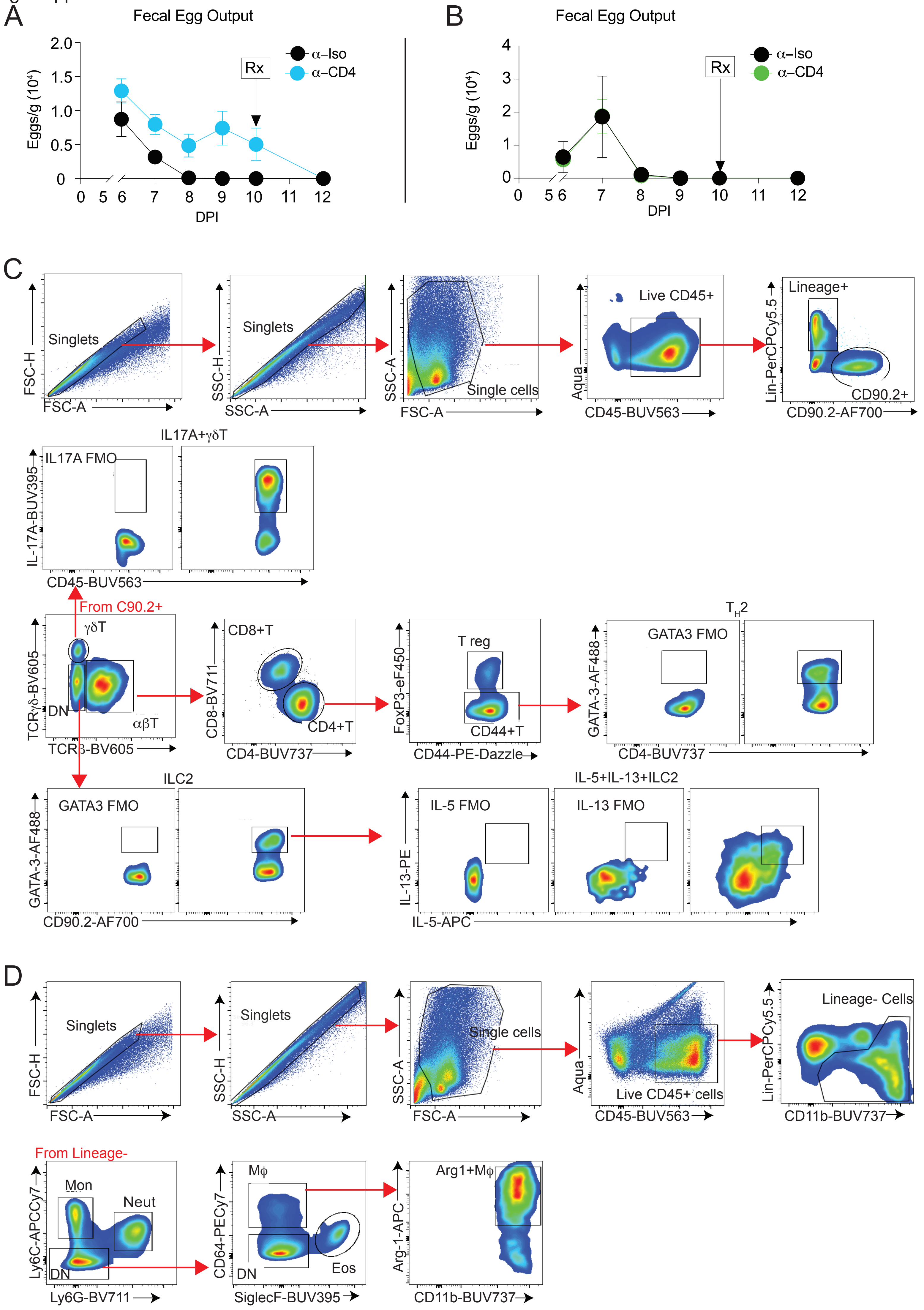

### Supplemental Figure 2

Fig. Supplemental 2

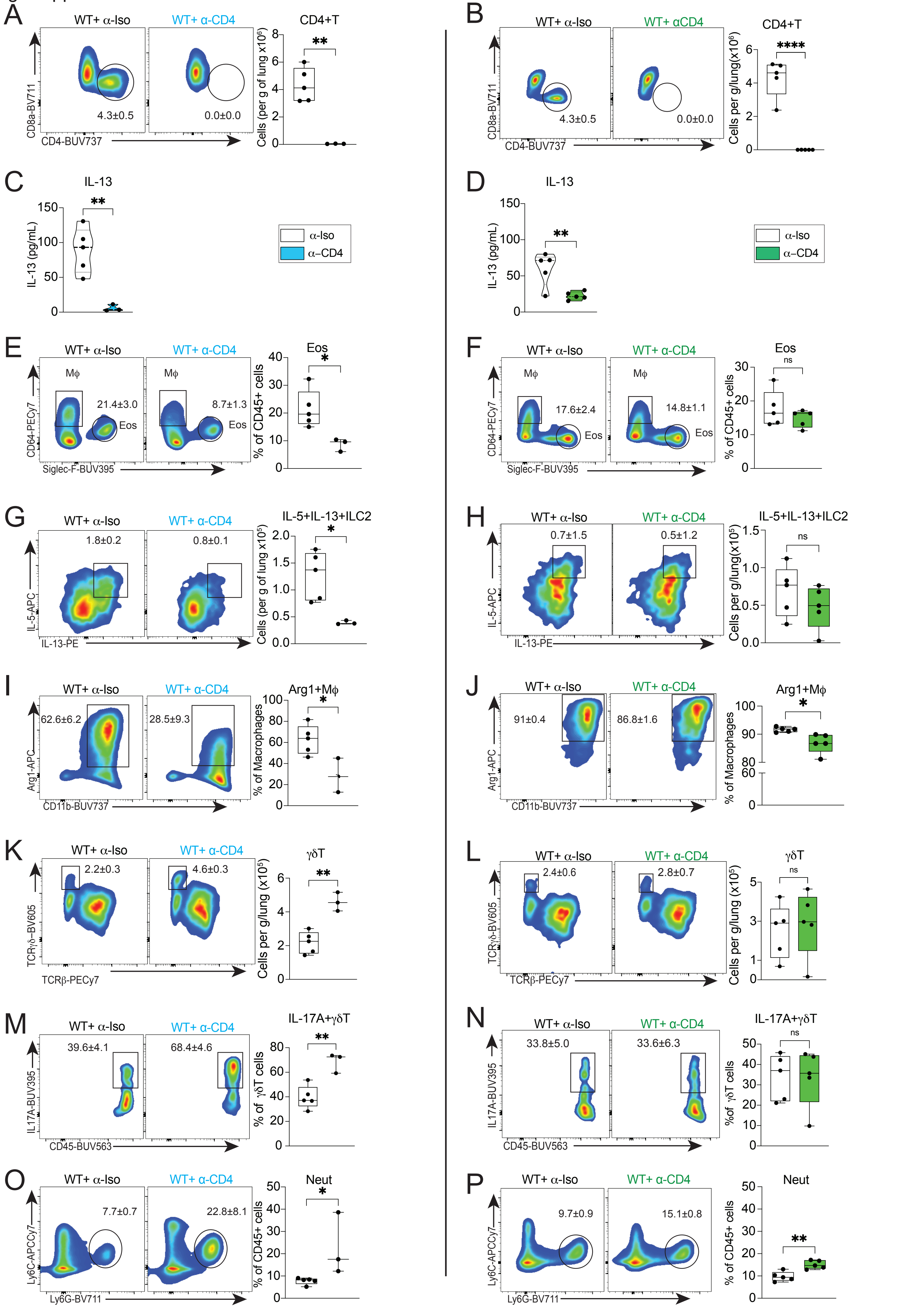

### Supplemental Figure 3

Fig. Supplemental 3

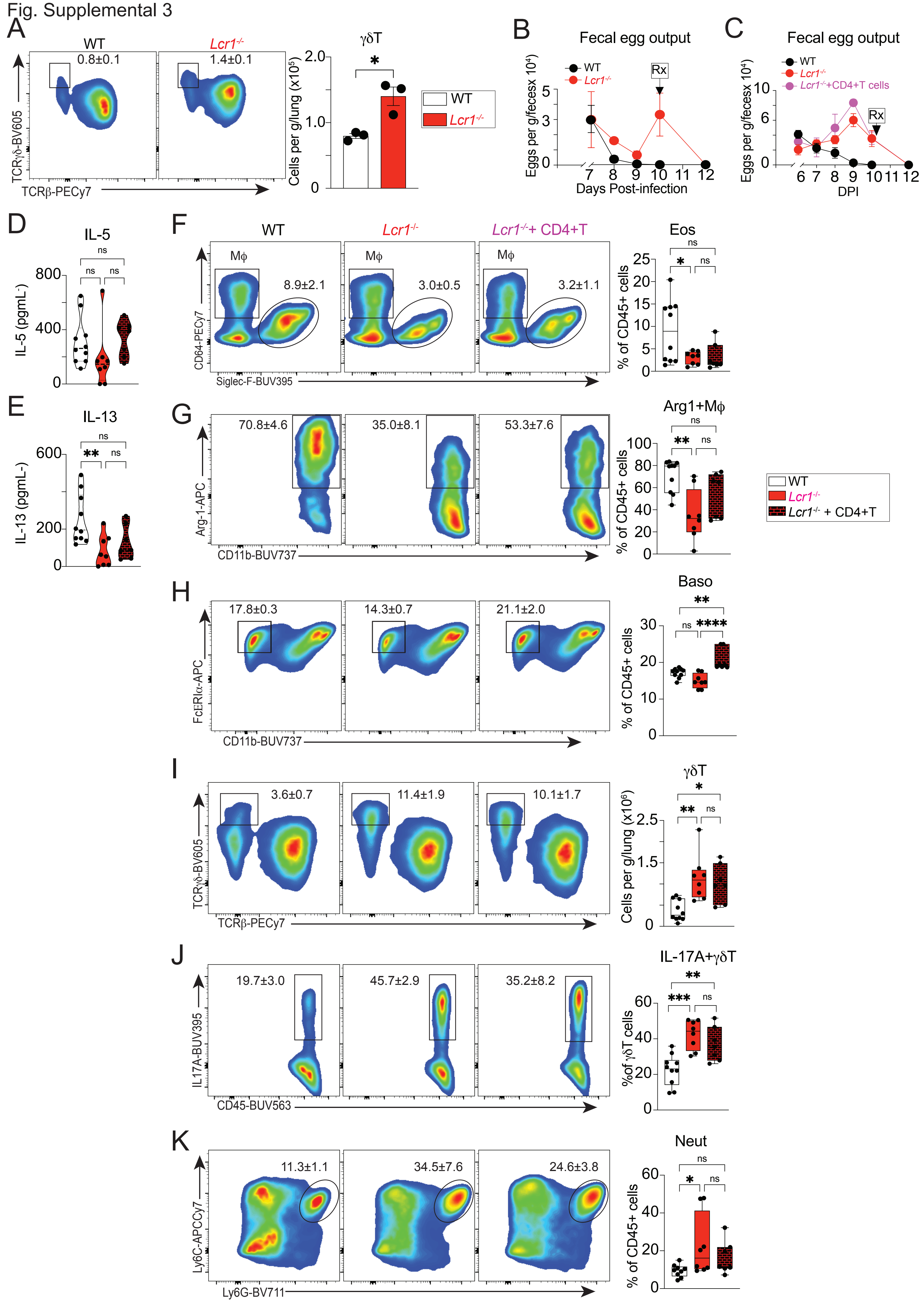

### Supplemental Figure 4

Fig. Supplemental 4

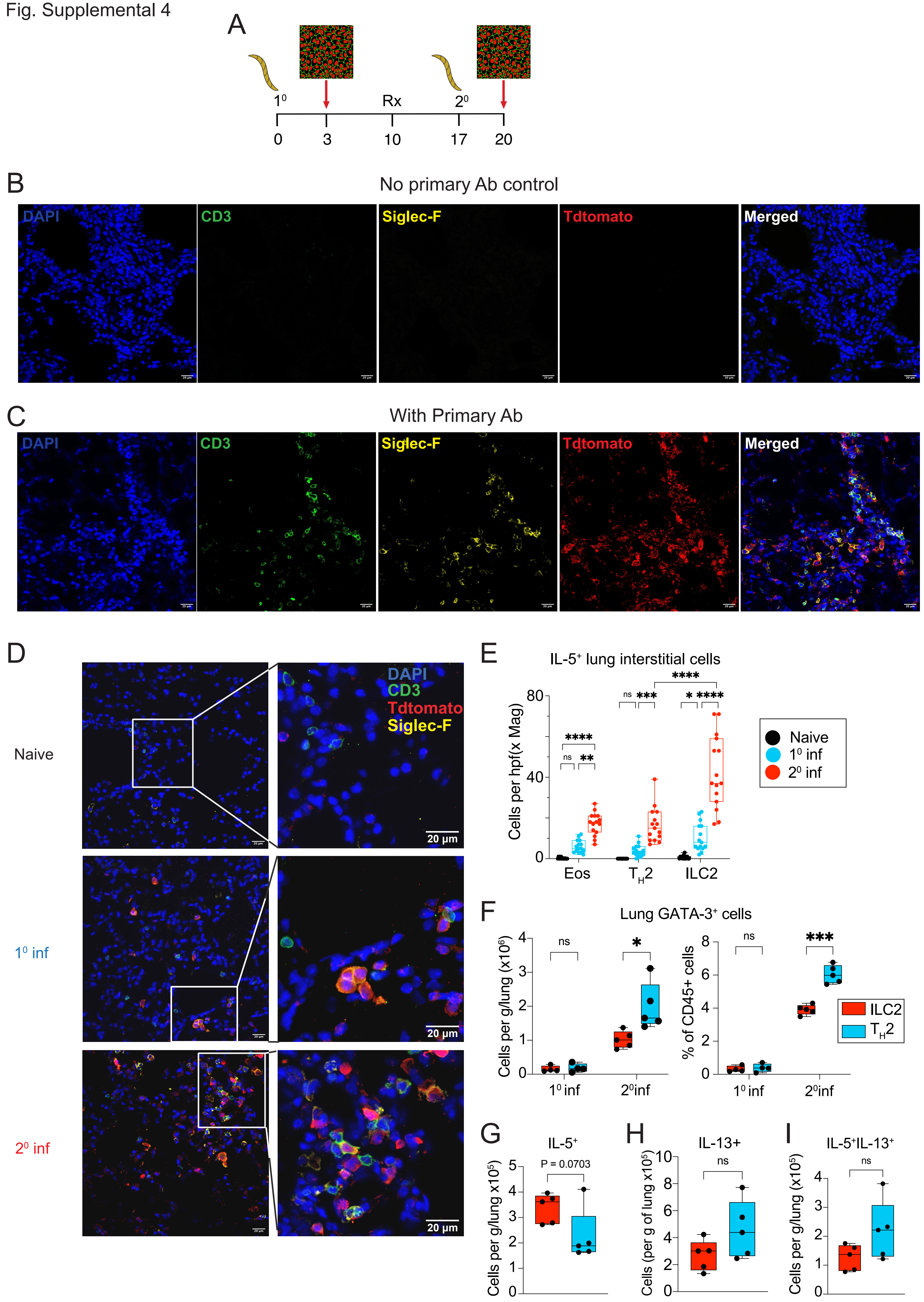

### Supplemental Figure 5

Fig. Supplemental 5

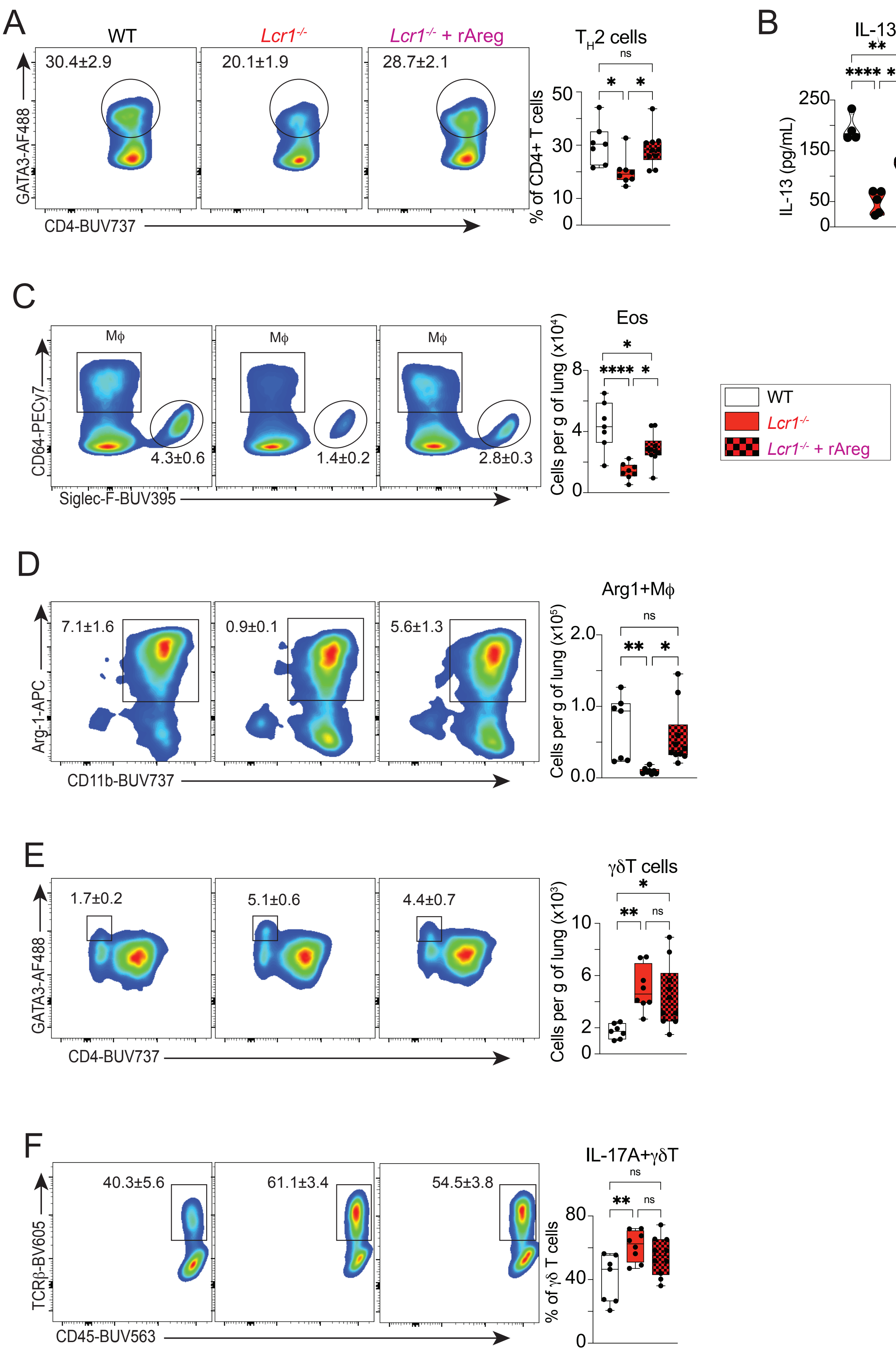

### Supplemental Figure 6

Fig. Supplemental 6

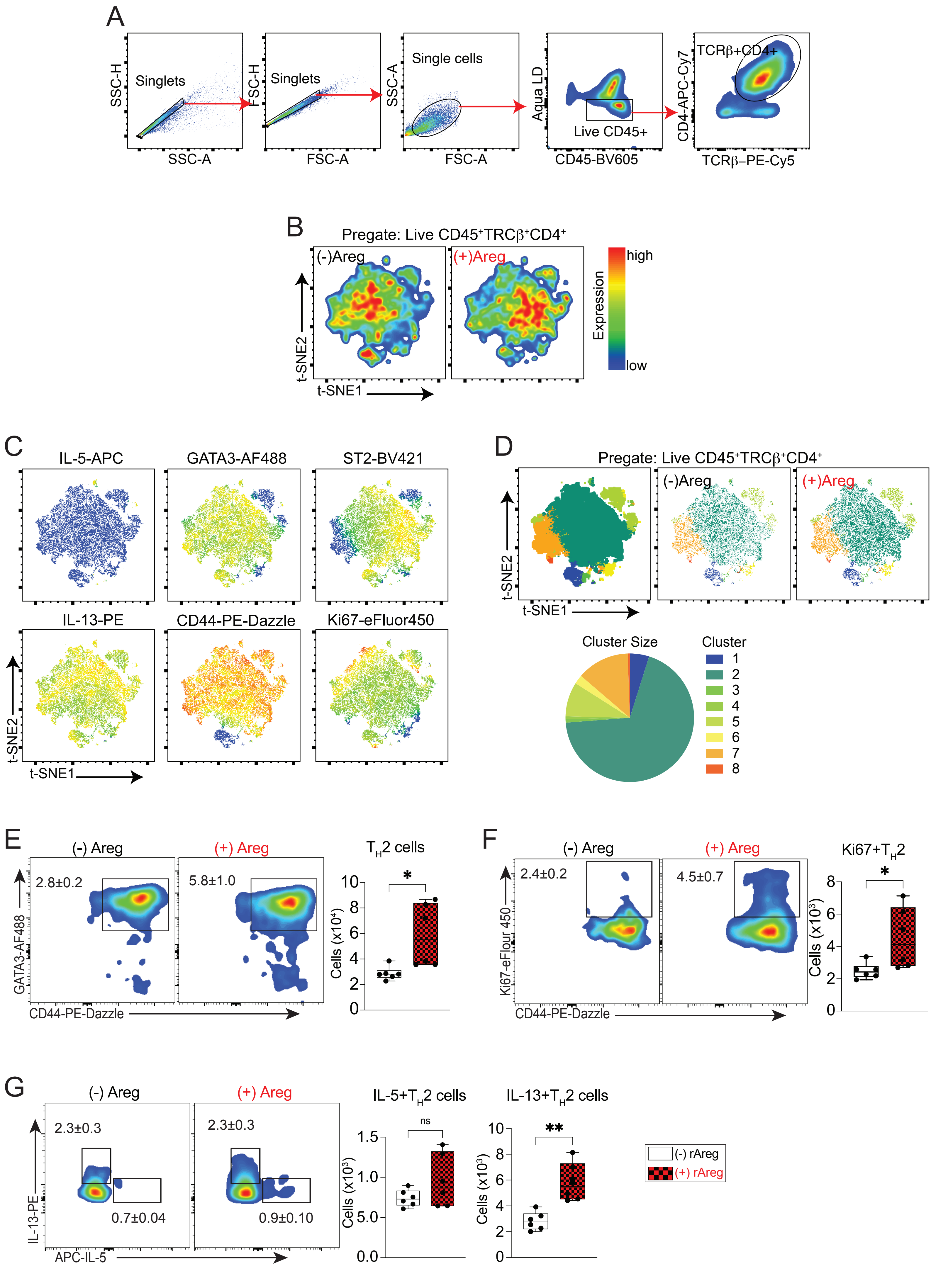

### Supplemental Figure 7

Fig. Supplemental 7

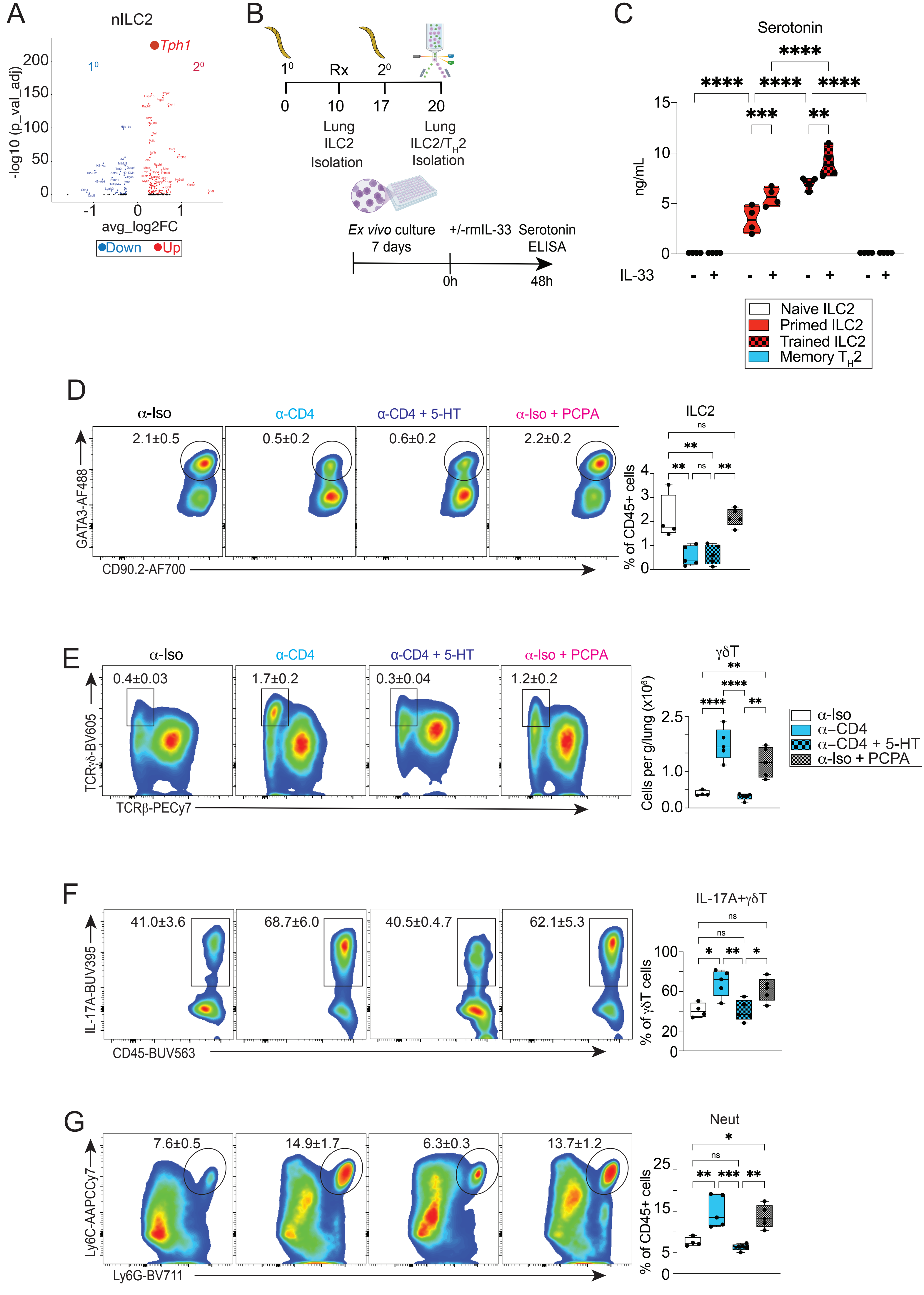
